# Curcumin and 5-bromoindole-3-carboxaldehyde inhibit the production of Quorum sensing-dependent virulence factors in *Pseudomonas aeruginosa*

**DOI:** 10.1101/2024.07.29.605599

**Authors:** Gabriel Martínez González, José Marcos Falcón-González, Rodolfo García Contreras, Juan Andres Alvarado-Salazar, Israel García Valdés, Nilda García Valdés, Denisse Alejandra Lugo Gutiérrez, Ingrid Lizeth Bustamante Martínez, Alonso Ruben Tescucano Alonso, Alejandra Ramirez-Villalva, Jorge A. Almeida

## Abstract

Curcumin and 5-bromoindole-3-carboxaldehyde were tested for their ability to inhibit quorum sensing-dependent virulence factors in *Pseudomonas aeruginosa* strains Pa01, Pa14, and ATCC 27853. The compounds were evaluated at curcumin (1, 5 μg/ml) and 5-bromoindole-3-carboxaldehyde (30, 50, 100 μg/ml). Virulence factors—exoprotease, elastase, pyocyanin, and alginate —were measured. Chemoinformatics properties were analyzed using Molinspiration, Molsoft, Osiris Property Explorer, pkCSM, and SwissADME. Results showed 5-bromoindole-3-carboxaldehyde significantly reduced pyocyanin in Pa01 (92.5% at 100 μg/ml), elastase (79.85% at 100 μg/ml), exoprotease (92.8% at 50 μg/ml), and alginate in ATCC 27853 (76.16% at 100 μg/ml). Curcumin effectively inhibited pyocyanin in ATCC 27853 (100% at 5 μg/ml), elastase in Pa14 (56.19% at 5 μg/ml), exoprotease in Pa01 (87.04% at 5 μg/ml), and alginate (48.26% at 5 μg/ml). Both compounds demonstrated strong inhibition of virulence factors, suggesting their potential as anti-virulence agents against *P. aeruginosa*, a highly resistant bacterium prevalent in nosocomial infections.

## Introduction

*Pseudomonas aeruginosa* is a Gram-negative aerobic rod-shaped bacterium that can be isolated from most environments, including soil, plants, and mammal tissue [1]. Has become an important cause of infection in humans and can be associated with significant morbidity and mortality [2]. Is a common cause of nosocomial infections, particularly pneumonia, infection in immunocompromised hosts, and in those with structural lung disease such as cystic fibrosis [3,4].

The World Health Organization (WHO) listed *P. aeruginosa* as one of three bacterial species in which there is a critical need for the development of new antibiotics to treat infections [5]. Importantly, *P. aeruginosa* is one of the MDR ESKAPE pathogens [6].

*P. aeruginosa* virulence regulation uses both chemical factors such as quorum sensing (QS) and mechanical cues such as surface association to modulate a large suite of virulence factors [7]. Essential function for QS in *P. aeruginosa* is to regulate the production of multiple virulence factors, such as extracellular proteases, iron chelators, efflux pump expression, biofilm development, swarming motility, phospholipases C, alginate, DNase and the response to host immunity [8,9].

Resistance of *Pseudomonas aeruginosa* to antibiotics is a major problem. Targeting virulence factors is an alternative option to avoid the emergence of resistance to antibiotics and treat infections that this pathogen causes [10]. QS system an attractive target for antibacterial therapy [11]. Inhibition of this system may cause the attenuation of virulence and protect against infection. In fact, an anti-QS approach has already shown promise in the battle against *P. aeruginosa* infections [12].

The use of QS inhibitors would be of particular interest in inhibiting bacterial pathogenicity and infections [13]. Turmeric (*Curcuma longa*) has traditionally been used as anti-inflammatory, antimicrobial and antifungal, antioxidant, anticancer, anti-inflammatory, antiobesity and antidepressant [14], but reports on its anti-QS properties are scarce [15]. Several studies reported that curcumin inhibits bacterial QS systems/biofilm formation and prevents bacterial adhesion to host receptors in various species, including *Staphylococcus aureus, Enterococcus faecalis, Escherichia coli, Streptococcus mutans, Listeria monocytogenes, Helicobacter pylori, Pseudomonas aeruginosa, Serratia marcescens, Aeromonas hydrophila* and *Acinetobacter baumannii* [16].

Another molecule whose reports indicate antibacterial activity are indoles, which, although they use bacteria as signalling, have shown important antibiofilm activity [17]. Functional alteration of indole carboxaldehydes may increase their effectiveness as quorum sensing inhibitors (QSIs). The bromination of carboxaldehydes represents an increase in their activity [18].

Given the background of the anti-quorum sensing activity of molecules such as curcumin and carboxaldehydes, the objective of this work is to evaluate in three strains of *Pseudomonas aeruginosa* (Pa01, Pa14 and ATCC 27853) the activity of curcumin and 5-bromoindole-3-carboxaldehyde on quorum-sensing-dependent virulence factors, such as alginate, pyocyanin, elastase and protease.

The proposal of new antibacterial treatments often requires the use of computer software that allows the calculation of molecular properties and the prediction of bioactivity, with which the cheminformatics properties, previously obtained, can be correlated with the viability of the new treatment proposal. Chemoinformatic properties have advantages such as predicting the site of action, probable route of administration, physicochemical, pharmacokinetic, pharmacodynamic behaviour and possible toxic effects. The use of simulation techniques and online servers allow us to know the behaviour of a compound to propose an effective and safe therapy.

## Methods and materials

### Preparation of strains and minimum inhibitory concentration

To carry out the tests, 3 different strains of *P. aeruginosa* were used; Pa01, Pa14 and ATCC 27853, all belonging to the microbiology laboratory of the University of Ixtlahuaca CUI. To activate these strains, they were reseeded in boxes of LB medium with 1.5% agar for at least 20 hours at 37 °.

The minimum inhibitory concentration (MIC) was developed using the microdilution method in a 96-well plate (Nunc), according to Table 8A of the Clinical and Laboratory Standards Institute (CLSI)[19,20]. To evaluate the minimum inhibitory concentration of each molecule, curcumin at concentrations of 1000, 500, 250, 125, 62, 31.25, and 15.5 µg/mL, and 5-bromoindole-3-carboxaldehyde at concentrations of 1000, 500, 250, 125, and 62.5 µg/mL dissolved in DMSO, were tested against strains Pa01, Pa14, and ATCC 27853. The inocula were adjusted to achieve 10⁸ CFU/mL of microorganisms. Subsequently, 10 µL of the corresponding inocula were added to the wells with Mueller Hinton Broth (MHB), obtaining a final concentration of 10⁵ CFU/mL [21]. The negative controls were: MHB without microorganisms, MHB with the *P. aeruginosa* strains, and MHB with DMSO and *P. aeruginosa* strains to confirm that it did not interfere with growth. The microplates were incubated at 35 ± 2°C for 20 h. The analysis of each molecule was performed in triplicate.

### Preparation of the strains and addition of evaluated molecules

To activate these strains: Pa01. Pa14 y ATCC 27853, they were reseeded in boxes of LB medium with 1.5% agar for at least 20 hours at 37 °. Subsequently, ON (overnight) cultures of each strain were inoculated in 50 ml flasks with 25 ml of LB medium, cultivated for 18 hours at 37 °. After the elapsed time, the growth of the cultures was measured at OD of 600 nm in a 1:10 ratio (dilution in LB).

Subsequently, new cultures of the strains were inoculated from the ON cultures with an OD of 600 nm. The inoculums of each strain of interest were added to 3 series of flasks (each with 8 flasks with 25 ml of LB medium) since three different strains were evaluated, different concentrations of each molecule were added to each series: For curcumin (Sigma-Aldrich), they evaluated concentrations of 1 µg/ml and 5 µg/ml, 5-bromoindole-3-carboxaldehyde (Sigma-Aldrich) 30, 50 and 100 µg/ml, both molecules dissolved in DMSO previously filtered with acrodiscs 0.22 µm. They were incubated at 37 ° with constant shaking for 24 hours. To maintain control, a culture that did not have any of the added molecules was used. After 24 hours, growth was measured at 600 nm of both the controls and the flasks with the evaluated molecules. As the molecules were dissolved in DMSO, negative controls were done with this solvent.

### Determination of Pyocyanin

From the mother flasks with ON, 1000 µl of each culture were taken. They were subsequently centrifuged at 13,000 rpm for 1.5 minutes. 800 µl of the supernatant were taken. Adding 420 µl of chloroform, it was vortexed for 1 minute at speed 10 and centrifuged at 13,000 rpm for 5 minutes. We transferred 300 µl of the bottom phase together with 800 µl of 0.1 N HCl. It was shaken for 1 minute at vortex speed 10 and centrifuged at 13,000 rpm for 5 minutes. It was determined quantitatively by absorbance, measured in a quartz cell, at 520 nm [22].

### Protease Determination

1000 µl were taken from the stock flasks with ON. The samples were centrifuged at 13000 rpm for 2 minutes. 875 µl of buffer with protease reagent and 125 µl of culture supernatant were used. It was vortexed for 1 minute. Samples were incubated at 37 ° at 200 rpm for 1 h. At the end of the incubation time, aliquots were taken and placed in 1% HN0_3_. Shaking with the help of vortex for 1 minute at speed 10, we then centrifuged at 13,000 rpm for 2 minutes. The supernatant was placed in the presence of 0.5% Na0H. We stir lightly with the help of the vortex. Transferring into quartz cells, the samples were read at 595 nm [23,24].

### Determination of alginate

1000 µl were taken from the stock flasks. 1 ml of the cell culture from the cultures was centrifuged and subsequently heated at 80° for 30 min. The resulting supernatant was centrifuged for 30 min at 13,000 rpm, the pellet was removed, and the alginate was precipitated with ice-cold 99% ethanol. The precipitated alginate was collected and dissolved in 1 ml of 0.9% sterile saline. A mixture of 118 µl of sample plus 1 ml of H_3_BO_3_/H_2_SO_4_ and 34 µl of carbazole is made. This mixture was heated for 30 min at 55°C and the absorbance (OD 530) was measured [25].

### Determination of Elastase

1000 µl were taken from the mother flasks, then centrifuged for 1.5 minutes at 13000 rpm and the supernatant was separated. Subsequently, 100 µl of the supernatant from each of the cultures was placed and 5 mg of elastin-Congo red from Sigma-Aldrich and 900 µl of pH 7.5 buffer were added. They were incubated for 2 h at 37 °, centrifuged at 13,500 rpm for 5 minutes, and absorbance was measured at 495 nm [26].

### Molecular docking

The 2D structure of curcumin and 5-bromoindole-3-carboxaldehyde were drawn with ChemSketch ACD/Labs and protonation states were calculated at pH 7.4 with MarvinSketch Chemaxon. The 3D structure was obtained and optimized to a semi-empirical PM3 level with Spartan 08 V1.2.0. The crystals of the LasR, PQS and RhlR proteins were obtained from the Protein Data Bank repository with the codes 3IX3, 4JVI and 8DQ0, respectively. The preparation of the proteins was carried out in Molegro Virtual Docker, the water molecules were removed, the druggable cavities were detected and the ligand-binding site was selected for molecular docking studies with a search sphere of 10 Å radius, grid of 0.30 Å, with the coordinates X: 10.54, Y: 4.32, Z: 20.98 for LasR, X: −34.71, Y: 56.02, Z: 9.17 for PQS and X: 116.01, Y: 96.69, Z: 137.48 for RhlR [27–29].

The validation of the molecular docking method was carried out using 12 methods based on the combinations of 4 scoring functions (MolDock Score, MolDock Score GRID, Plant Score and Plant Score GRID) and 3 search algorithms (MolDock Optimizer, MolDock SE and Iterated Simplex). In each method, 10 runs were performed, a maximum of 1500 interactions, and an initial population of 50 poses [27, 28]. The method for performing molecular docking, which consists of the scoring function and search algorithm, was chosen considering an RMSD value ≤ 2.0 as inclusion criteria [30], obtained for coupled ligand with respect to the position of the co-crystallized ligand. The binding energy was obtained in the online server PROtein binDIng enerGy prediction (PRODIGY) [31], and the interactions were visualized with the BIOVIA Discovery Studio program [32].

### Molecular dynamics simulations

To evaluate the dynamic stability and conformational flexibility of the protein-ligand complexes from molecular docking under physiological conditions, Molecular Dynamics (MD) simulations were performed using GROMACS 2024-3 software [33,34]. The force field used within MD considered all atoms and it was consistent with the CHARMM36 parameter set [35]. For proteins LasR, PQS and RHIR, the parametrization for this force field is straightforward and the topology is automatically generated by GROMACS. In the case of curcumin and 5-bromoindole-3-carboxaldehyde the generation of topologies and the assignment of parameters were conducted by the CGenFF Web App by SilcsBio [36].

After topologies and force field parameters were generated and validated, each of the protein-ligand systems were placed in the center of a simulation box and subsequently solvated with liquid water using the TIP3P potential [37]. Also, Cl^−^ and Na^+^ ions were added in a proportion that would neutralize the overall charge of the system and obtain a final salt concentration of 100 mM. Both the energy minimization to avoid spurious contacts and the equilibration of systems were carried out according to the protocol and parameters set outlined in [38].

For production and obtaining observables, MD simulations were performed in the isothermal-isobaric ensemble (NPT). The Nose-Hoover thermostat and Parrinello-Rahman barostat were used to keep a constant temperature of 310 K at a pressure of 1 bar, respectively [39]. The time interval of Δt = 0.002 ps was established to integrate the equations of motion. Then, 200 ns trajectories were generated for each system. Hydrogen bonding interactions, root mean square deviation (RMSD), root mean square fluctuation (RMSF) and binding energy were calculated from MD simulations.

To obtain an appropriate estimate of the binding energy for the protein-ligand complex, we used the Molecular Mechanics Poisson-Boltzmann Surface Area (MM/PBSA) method [40]. In broad terms, MM/PBSA takes into account the gas phase free energy contribution (*ΔE_MM_*), the solvation free energy (*ΔE_solv_*) and the conformational entropy (*-T ΔS*):

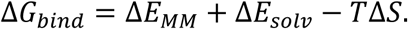

In this framework, the gas phase free energy considers both electrostatic and van der Waals contributions (*ΔE_MM_*= *ΔE_ele_* + *ΔE_vdw_*), while the solvation free energy comprises both polar and non-polar contributions (*ΔE_solv_*= *ΔE_pol_* + *ΔE_non-pol_*). The entropy term (*-T ΔS*) was not considered because its calculation is computationally demanding and the results are not reliable [41]. MMPBSA calculations also exploit the trajectories generated from MD simulations.

### Statistical analysis

Two-way analysis of variance (ANOVA) with replication was performed to evaluate the effect of different treatments on bacterial strains Pa01, Pa14, and ATCC27853. The treatments included 5-bromoindole-3-carboxaldehyde at three different concentrations (30 µg/ml, 50 µg/ml, and 100 µg/ml), and curcumin at two different concentrations (1 µg/ml and 5 µg/ml). Both the ANOVA analysis and the calculation of standard deviation (SD) were performed using IBM SPSS Statistics for Windows, version 21.0 (IBM Corp., Armonk, NY).

## Results

### Experimental evaluation

In the present study were the effectiveness of two compounds, 5-bromoindole-3-carboxaldehyde and curcumin at different concentrations, in the inhibition of pyocyanin, alginate, elastase and protease quorum-dependent virulence factors, and minimal inhibitory concentration (MIC) in three *Pseudomonas aeruginosa* strains was evaluated: Pa01, Pa14 and ATCC 27853.

### Minimal inhibitory concentration (MIC)

The MIC values evaluated for curcumin showed no inhibitory activity in the 15.5–250 µg/mL range. Growth inhibition was evident only at concentrations ≥500 µg/mL, and up to 1000 µg/mL. In contrast, 5-bromoindole-3-carboxaldehyde showed significant inhibitory activity in all strains starting at a concentration of 125 µg/mL, without a progressive increase in concentration-dependent inhibition Table 1.

**Table 1.**
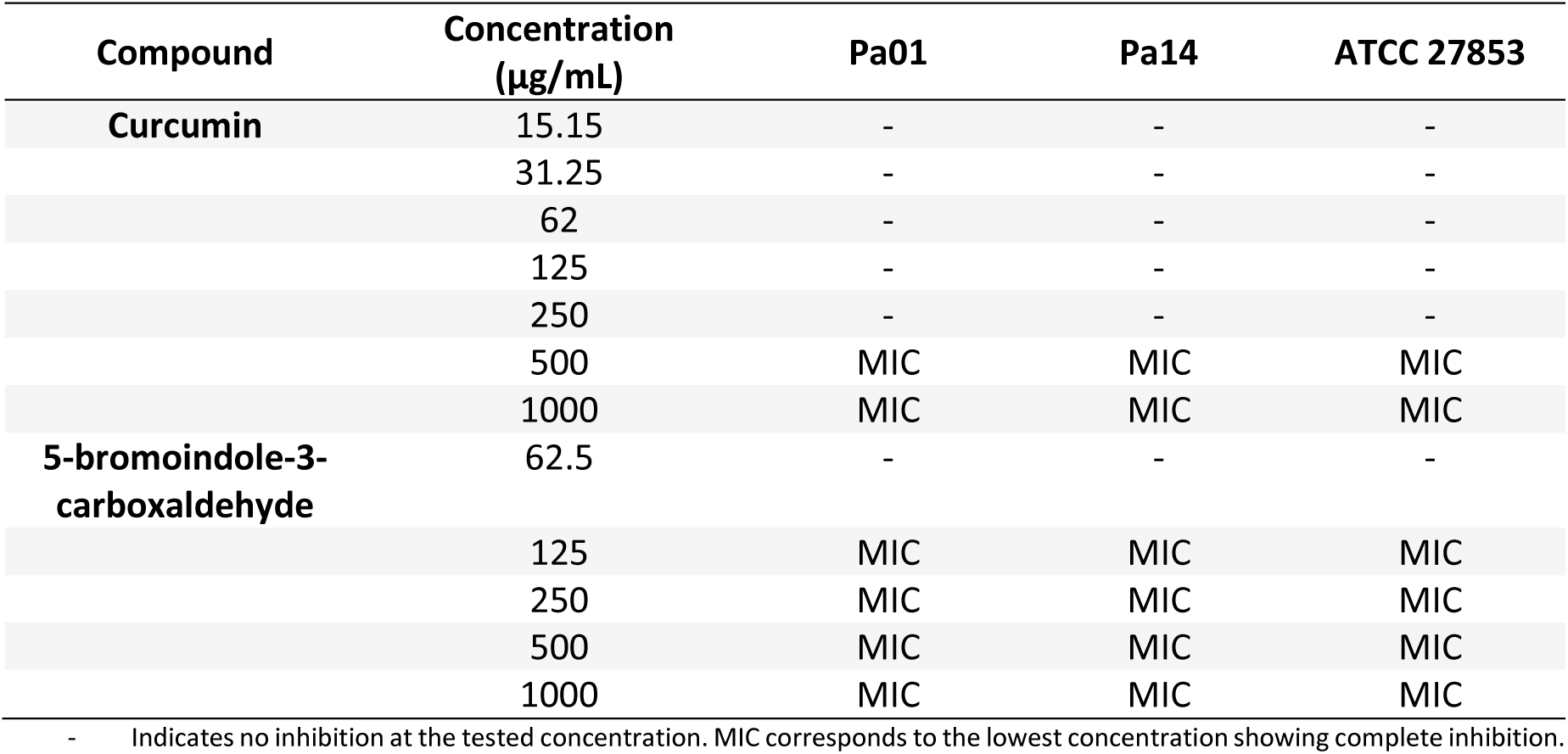
Minimum inhibitory concentrations (MIC) of curcumin and 5-bromoindole-3-carboxaldehyde against three *Pseudomonas aeruginosa* strains.

#### Pyocyanin

Bromoindole: Strain Pa01 showed significant inhibition at concentrations of 30 µg/ml (85.42%), 50 µg/ml (87.77%) and 100 µg/ml (92.5%). The Pa14 strain also presented a reduction, although smaller, with values of 60.3%, 61.67% and 67.65% at the same concentrations, respectively. Strain ATCC 27853 showed lower inhibition at 30 µg/ml (8.7%) and 50 µg/ml (13.05%), but higher inhibition at 100 µg/ml (72.47%). Regarding the Pa14 strain at the same concentration. Although the Pa01 strain was the one that produced the highest concentration of pyocyanin in its control, it was the strain in which the molecule had the highest activity at all the concentrations studied. On the other hand, the control production of the Pa14 strains and the ATCC 27853 strain varies by 0.001, showing greater activity in the Pa14 strain at concentrations of 30 and 50 µg/ml, but not for the concentration of 100 µg/ml where it is seen. greater inhibition in strain ATCC 27853.

When curcumin was added, strain Pa01 showed an inhibition of 60.68% at 1 µg/ml and 90.63% at 5 µg/ml. The Pa14 strain presented inhibitions of 19.12% at 1 µg/ml and 55.89% at 5 µg/ml. The ATCC 27853 strain showed inhibitions of 81.16% at 1 µg/ml and 100% at 5 µg/ml. Although total inhibition is achieved in strain ATCC 27853, it is the one that showed the lowest pyocyanin production, however, it is the strain with the highest activity, followed by Pa01 and Pa14 respectively. Graph 1.

**Graph 1.**
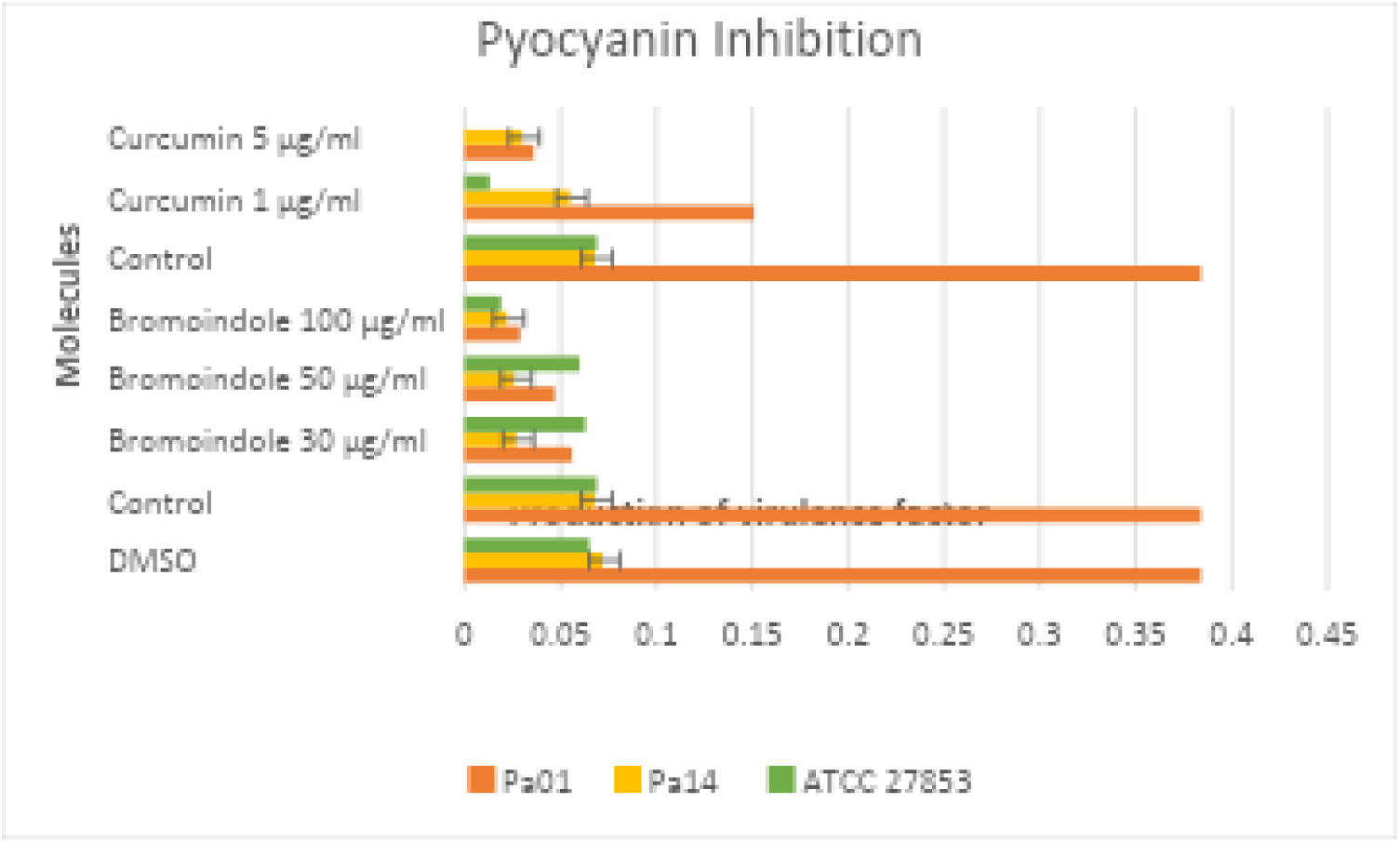
Pyocyanin inhibition in the presence of different concentrations of curcumin and 5-bromoindole-3-carboxaldehyde

The two-factor analysis of variance found no significant differences in the inhibition of pyocyanin production between the different treatments with 5-bromoindole-3-carboxaldehyde and Curcumin (F (5, 5) = 1.448, p = 0.347) and between the different bacterial strains evaluated (F (1, 5) = 0.003, p = 0.961).

#### Elastase

Bromoindole: For strain Pa01, a significant inhibition was observed with values of 69.77% at 30 µg/ml, 70.64% at 50 µg/ml and 79.85% at 100 µg/ml. The Pa14 strain showed a variable behaviour since when adding 30 µg/ml no inhibition was observed, on the contrary it seems that it favoured the expression of the virulence factor with 21.11% more compared to the control, an inhibition of 30.42% was shown at 50 µg/ml and 32.99% at 100 µg/ml. The ATCC 27853 strain showed an initial inhibition at 100 µg/ml (31.35%) since, as happened with the Pa14 strain, at concentrations of 30 and 50, greater production of the virulence factor was shown, which suggests that at low concentrations it does not It inhibits but it does stimulate production. Therefore, the molecule showed a high inhibitory effect at all concentrations studied in the Pa01 strain, for the Pa14 strain it is the second in terms of activity and in which it showed the least activity was in the ATCC 27853 strain.

The Pa01 strain presented inhibitions when adding curcumin of 3.6% at 1 µg/ml and 32.08% at 5 µg/ml. Strain Pa14 showed minor inhibitions of 5.93% at 1 µg/ml and 56.19% at 5 µg/ml. The ATCC 27853 strain presented inhibitions of 5.65% at 1 µg/ml and 18.5% at 5 µg/ml. The molecule at 1 µg/ml presented the least inhibition in the Pa01 strain, and greater activity at both concentrations in the Pa14 strain. Graph 2.

**Graph 2.**
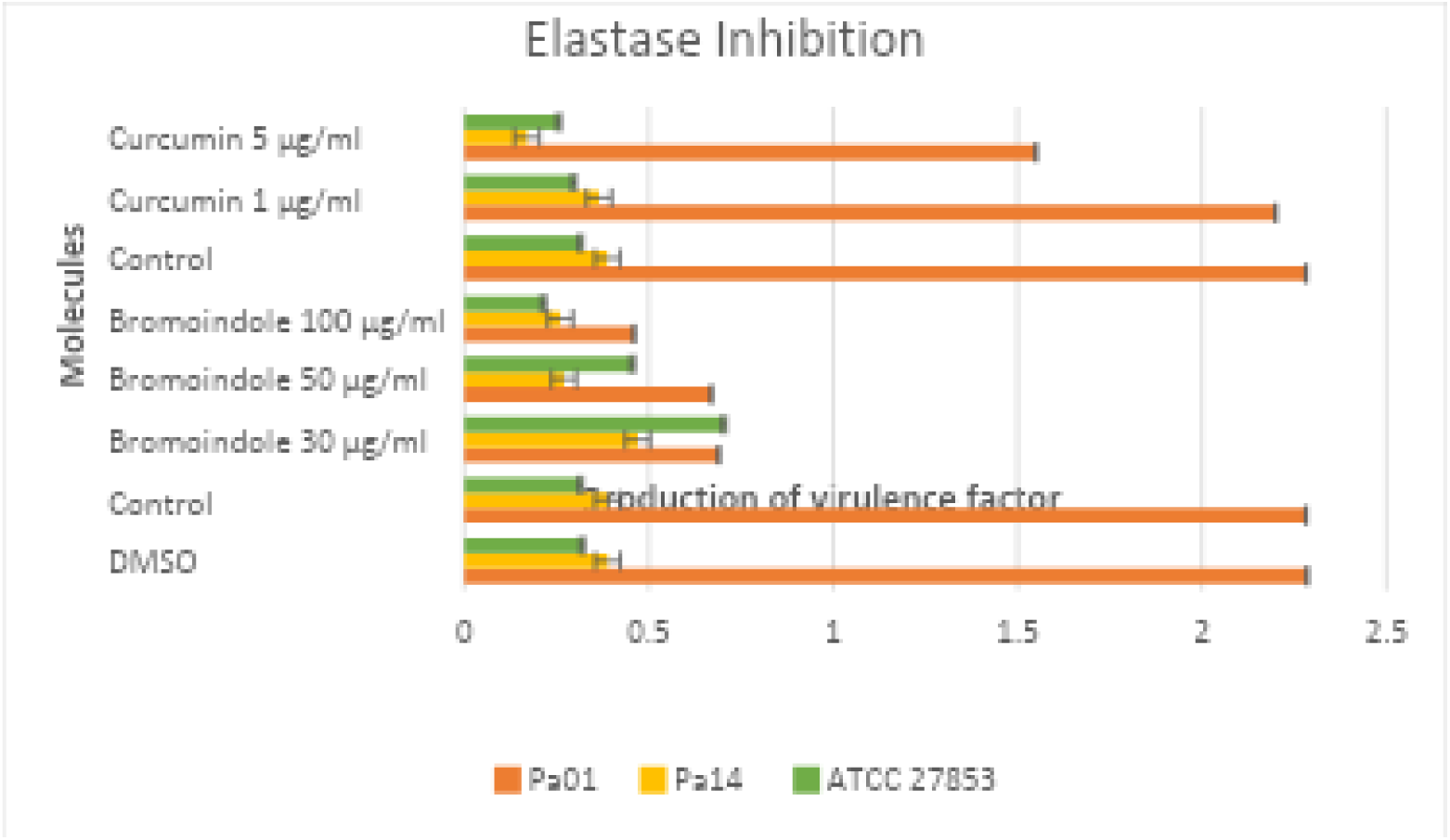
Elastase inhibition in the presence of different concentrations of curcumin and 5-bromoindole-3-carboxaldehyde

The two-factor analysis of variance did not find significant differences in the inhibition of elastase production between the different treatments with 5-bromoindole-3-carboxaldehyde and Curcumin (F (5, 5) = 3.844, p = 0.083) and between the different strains evaluated (F (1, 5) = 1.076, p = 0.347). The variability due to error was relatively low (0.0092), indicating that most of the total variability observed in the data can be explained by the molecules, although not significantly in terms of the strains.

#### Protease

Bromoindole: Inhibition in strain Pa01 was 92.51% at 30 µg/ml, 92.8% at 50 µg/ml and 89.92% at 100 µg/ml. Being the strain where at low concentrations of 30 µg/ml and 50 µg/ml a considerable inhibition is seen, however at 100 µg/m no greater inhibition is seen, on the contrary it seems that it decreases by about 3% compared to concentration No. 2. Strain Pa14 showed variable inhibition, with an initial increase at 30 µg/ml (66.6%), a reduction at 50 µg/ml (24.25%), and almost complete inhibition at 100 µg/ml (96.22%). The ATCC 27853 strain presented an inhibition of 85.56% at 50 µg/ml and 97.48% at 100 µg/ml, although at 30 µg/ml it also showed an increase in the expression of the virulence factor. However, for this virulence factor at concentrations of 30 and 50 µg/ml it is more effective in the Pa01 strain, while at the concentration of 100 µg/ml in the strain that showed the greatest activity in ATCC 27853 and in the one that showed the least activity at the same concentration was Pa01.

For curcumin, the inhibition in strain Pa01 was 79.83% at 1 µg/ml and 87.04% at 5 µg/ml. The Pa14 strain presented inhibitions of 1.52% at 1 µg/ml and 38.64% at 5 µg/ml. The ATCC 27853 strain showed inhibitions of 41.51% at 1 µg/ml and 71.12% at 5 µg/ml. The strain that showed the greatest inhibition of this virulence factor is Pa01, followed by ATCC 27853 and the one that showed the least activity against the molecule was Pa14. Graph 3.

**Graph 3.**
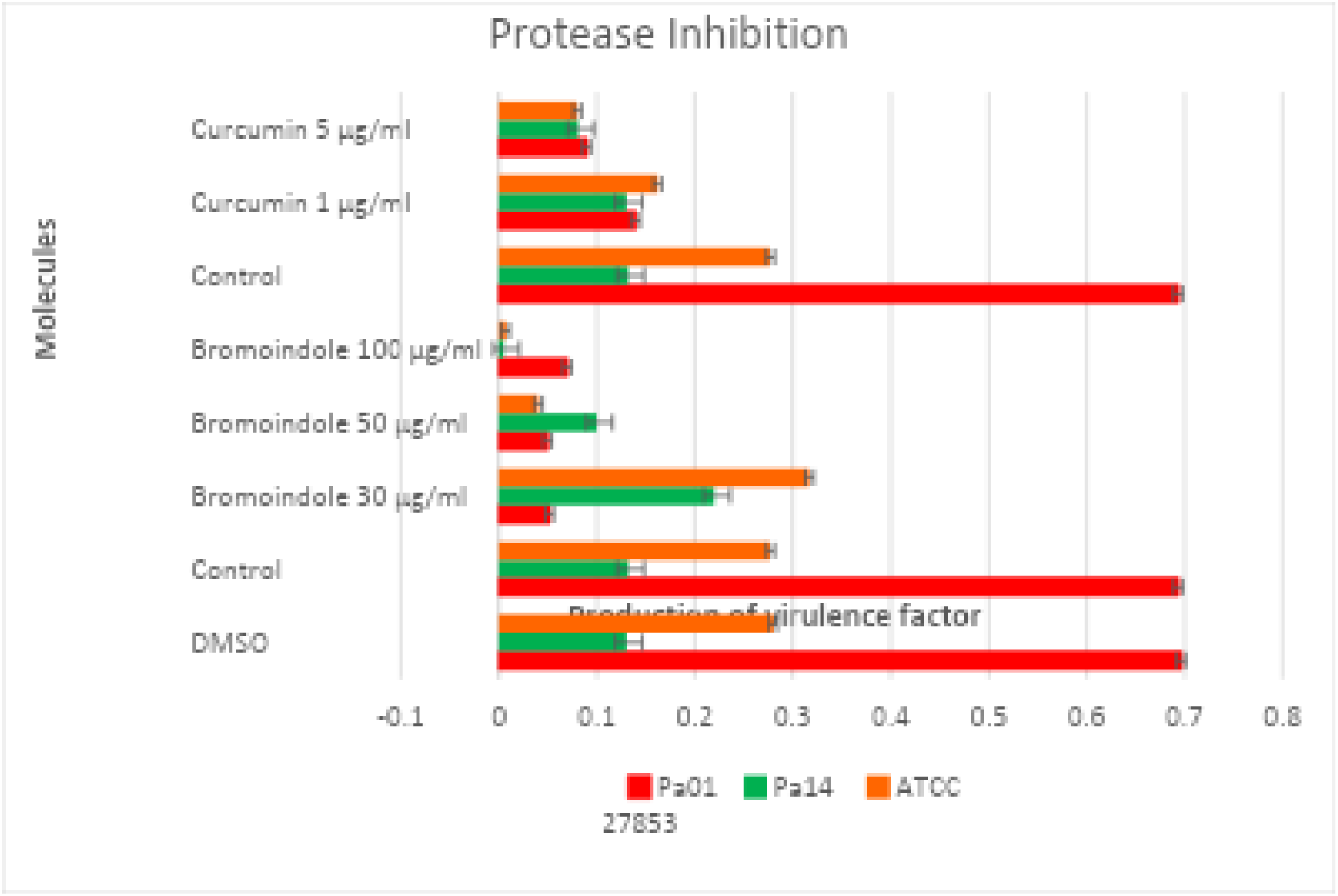
Protease inhibition in the presence of different concentrations of curcumin and 5-bromoindole-3-carboxaldehyde

The analysis of variance showed significant differences between the applied treatments in terms of protease activity (F (5, 5) = 6.786, p = 0.028). This suggests that at least one of the treatments (at different concentrations of Indole and Curcumin) has a significant effect in reducing protease activity compared to the control. The analysis did not indicate significant differences in the response to treatment between the bacterial strains Pa01, Pa14 and ATCC 27853(F (1, 5) = 1.408, p = 0.289). This implies that the strains do not respond differentially to the molecules tested. The variability due to error was relatively low (0.0027), indicating that most of the total variability observed in the data can be explained by the molecules.

#### Alginate

Bromoindole: Strain Pa01 showed an inhibition of 35.87% at 30 µg/ml, 48.4% at 50 µg/ml and 61.4% at 100 µg/ml. The Pa14 strain presented inhibitions of 16.27% at 30 µg/ml, 18.17% at 50 µg/ml and 40.67% at 100 µg/ml. Strain ATCC 27853 showed an initial increase in alginate production at 30 µg/ml (12.22%), but a significant inhibition at 50 µg/ml (52.78%) and 100 µg/ml (76.16%). The molecule was more active in strain Pa01 and ATCC 27853 at concentrations of 50 and 100 µg/ml, where the greatest activity is seen in the latter strain. For Pa14 it showed the least activity, but with significant inhibition.

With curcumin, strain Pa01 showed inhibitions of 37.91% at 1 µg/mL and 48.26% at 5 µg/ml. The Pa14 strain presented inhibitions of 32.3% at 1 µg/ml and 41.63% at 5 µg/ml. The ATCC 27853 strain showed inhibitions of 2.78% at 1 µg/ml and 15% at 5 µg/ml. Graph 4.

**Graph 4.**
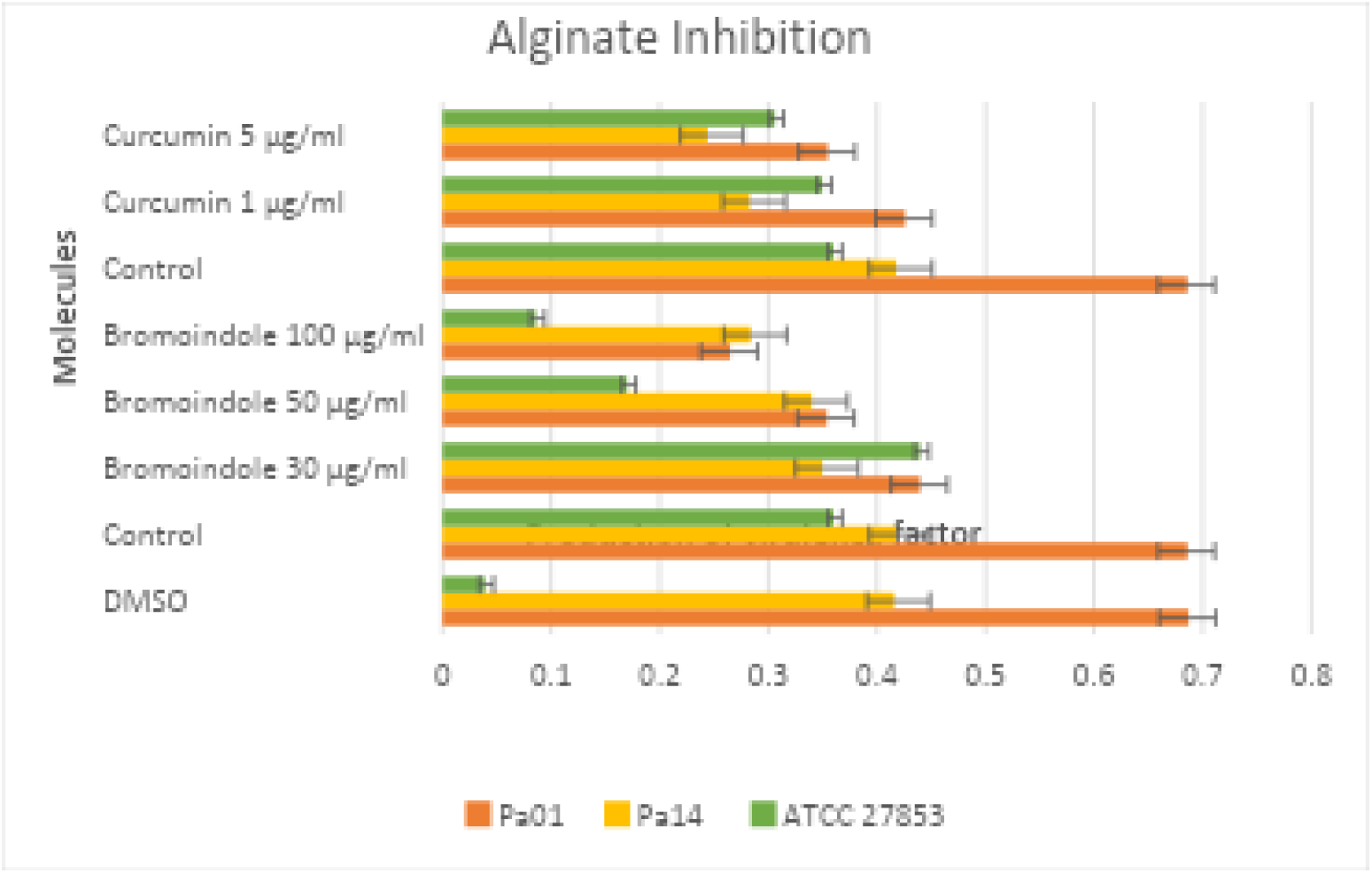
Alginate inhibition in the presence of different concentrations of curcumin and 5-bromoindole-3-carboxaldehyde

The analysis of variance did not find significant differences in alginate activity between the different treatments with 5-bromoindole-3-carboxaldehyde and Curcumin (F (5, 5) = 1.637, p = 0.301). Likewise, no significant differences were observed in the response to treatment between the strains evaluated (F (1, 5) = 0.442, p = 0.536).

Table 2 shows the results in percentage and inhibition concentration of the virulence factors; pyocyanin, elastase, protease and alginate studied with bromoindole. And table 3 shows the results in percentage and inhibition concentration of the virulence factors with curcumin.

**Table 2.** Inhibition percentages of 5-bromoindole-3-carboxaldehyde on Quorum-dependent virulence factors.

| Virulence factors |  |  |  |  |  |  |  |  |  |  |  |  |  |  |  |  |
| --- | --- | --- | --- | --- | --- | --- | --- | --- | --- | --- | --- | --- | --- | --- | --- | --- |
| Strain<br>/ □ | Pyocyanin |  |  |  | Elastase |  |  |  | Protease |  |  |  | Alginate |  |  |  |
|  | Control | 30<br>µg/ml | 50<br>µg/ml | 100<br>µg/ml | Control | 30<br>µg/ml | 50<br>µg/ml | 100<br>µg/ml | Control | 30<br>µg/ml | 50<br>µg/ml | 100<br>µg/ml | Control | 30<br>µg/ml | 50<br>µg/ml | 100<br>µg/ml |
| Pa01 | 0<br>(0.384) | 85.42<br>(0.56) | 87.77<br>(0.047) | 92.5<br>(0.029) | 0<br>(2.282) | 69.77<br>(0.69) | 70.64<br>(0.67) | 79.85<br>(0.46) | 0<br>(0.694) | 92.51<br>(0.052) | 92.8<br>(0.05) | 89.92<br>(0.07) | 0<br>(0.686) | 35.87<br>(0.44) | 48.4<br>(0.354) | 61.4<br>(0.265) |
| Pa14 | 0<br>(0.068) | 60.3<br>(0.027) | 61.67<br>(0.026) | 67.65<br>(0.022) | 0<br>(0.388) | +21.11<br>(0.47) | 30.42<br>(0.27) | 32.99<br>(0.26) | 0<br>(0.132) | +66.6<br>(0.22) | 24.25<br>(0.1) | 96.22<br>(0.005) | 0<br>(0.418) | 16.27<br>(0.35) | 18.17<br>(0.34) | 40.67<br>(0.284) |
| ATCC<br>27853 | 0<br>(0.069) | 8.7<br>(0.063) | 13.05<br>(0.06) | 72.47<br>(0.019) | 0<br>(0.319) | +121.1<br>(0.708) | +44.22<br>(0.46) | 31.35<br>(0.219) | 0<br>(0.277) | +14.44<br>(0.317) | 85.56<br>(0.04) | 97.48<br>(0.007) | 0<br>(0.36) | +12.22<br>(0.44) | 52.78<br>(0.17) | 76.16<br>(0.086) |

**Table 3.** Inhibition percentages of curcumin on Quorum-dependent virulence factors.

| Strain/ [] | Virulence factors |  |  |  |  |  |  |  |  |  |  |  |
| --- | --- | --- | --- | --- | --- | --- | --- | --- | --- | --- | --- | --- |
|  | Pyocyanin |  |  | Elastase |  |  | Protease |  |  | Alginate |  |  |
|  | Control | 1<br>µg/ml | 5<br>µg/ml | Control | 1<br>µg/ml | 5<br>µg/ml | Control | 1<br>µg/ml | 5<br>µg/ml | Control | 1<br>µg/ml | 5<br>µg/ml |
| **Pa01** | 0<br>(0.384) | 60.68<br>(0.151) | 90.63<br>(0.036) | 0<br>(2.282) | 3.6<br>(2.2) | 32.08<br>(1.55) | 0<br>(0.695) | 79.83<br>(0.14) | 87.04<br>(0.09) | 0<br>(0.686) | 37.91<br>(0.426) | 48.26<br>(0.355) |
| **Pa14** | 0<br>(0.068) | 19.12<br>(0.055) | 55.89<br>(0.03) | 0<br>(0.388) | 5.93<br>(0.365) | 56.19<br>(0.17) | 0<br>(0.132) | 1.52<br>(0.13) | 38.64<br>(0.081) | 0<br>(0.418) | 32.3<br>(0.283) | 41.63<br>(0.244) |
| **ATCC**<br>**27853** | 0<br>(0.069) | 81.16<br>(0.013) | 100<br>(0) | 0<br>(0.319) | 5.65<br>(0.301) | 18.5<br>(0.26) | 0<br>(0.277) | 41.51<br>(0.162) | 71.12<br>(0.08) | 0<br>(0.36) | 2.78<br>(0.35) | 15<br>(0.306) |

### Computational Studies

Molecular and cheminformatics properties were obtained using the Molinspiration, Molsoft, ChemSketch, swissADME and Osiris Property Explorer simulators. The property values of curcumin and 5-bromoindole-3-carboxaldehyde (Figure 1) are summarized in Table 3. Lipinski’s rules indicate that a compound must meet the following criteria to have good bioavailability and to be preferably administered orally: a) LogP value <5; b) TPSA <140 Å; c) molecular weight <500 Da; d) Hydrogen bond donors (-NH, -OH) <5; e) Hydrogen bond acceptors (N, O) <10 [42]; The Lipinski’s rule of 5 aids in differentiating between drug-like and non-druglike features and forecasts a high likelihood of success or failure due to a molecule’s drug-likeliness [43]. As can be seen in the table, curcumin and 5-bromoindole-3-carboxaldehyde do not present violations of Lipinsky’s rules, this would indicate that both compounds are candidates for oral administration, which is an advantage in antibacterial therapy.

**Figure 1.**
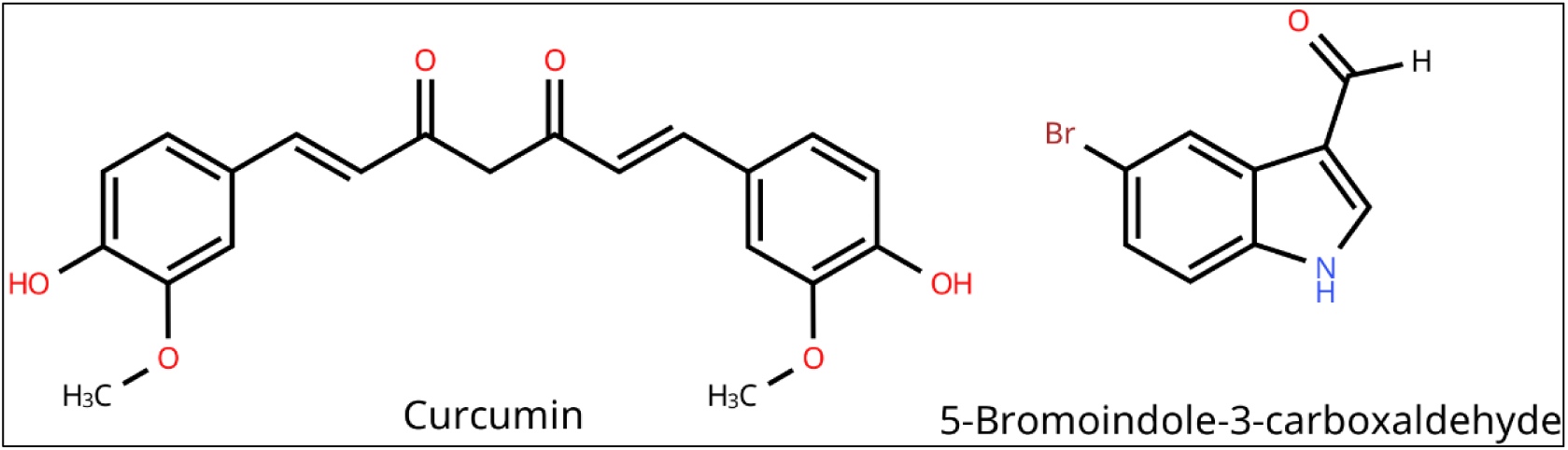
Structure of curcumin and carboxaldehyde

The characteristics to highlight include solubility, the Log P is relevant to drug discovery, as they are used to describe lipophilicity which influences drug-target, a relatively high lipophilicity results in reduced aqueous solubility and increased likelihood of metabolic instability [44], as seen in Table 4, curcumin has a LogP of 2.30 and 5-BIC of 2.66, therefore, both compounds are water-soluble, the PSA indicates the polar area surface of a molecule and the ease with which they can penetrate the cell, for which it must be less than 140 Å, curcumin has a PSA of 93.07 and 5-BIC of 32.86, this is indicative that the molecules could easily penetrate the cell membrane to carry out an effect on the cellular organelles. The presence of a hydrogen bond donor generally implies that a hydrogen bond acceptor is also present, these properties indicate electrostatic interactions resulting from aligned hydrogen bond donors and acceptors, which is useful in the recognition of molecular targets [45]. The topological descriptors of hydrophobicity (LogP), molar refractivity (MR), polarizability (P) and polar surface area (PSA) play an important role when estimating their activity and are generally related to the drug-active site interaction. Compounds with a water-soluble LogP usually present greater biological activity. Likewise, molar refractivity values close to 70 coincide with an increase in activity. Low polarizability (less than 50) indicates that the compound is specific, that is, it does not have the capacity to bind to several ligands, but rather its specificity for a receptor or target site increases. The data in Table 4 indicate that the presence of polar atoms in the molecule (but not polarizable) is necessary for these to be active. These results strengthen the proposal to be administered orally [46].

**Table 4.** Molecular and chemoinformatic properties of curcumin and 5-bromoindole-3-carboxaldehyde.

| *Properties* | *Curcumin* | *5-bromoindole-3-carboxaldehyde* |
| --- | --- | --- |
| **MF** | C<sub>21</sub>H<sub>20</sub>O<sub>6</sub> | C<sub>9</sub>H<sub>6</sub>BrNO |
| **MW** | 368.3799 | 224.05404 |
| **No. HBA** | 6 | 2 |
| **No. HBD** | 2 | 1 |
| **PSA** | 93.07 | 32.86 |
| **LogP** | 2.30 | 2.66 |
| **δ** | 1.279 ± 0.06 g/cm<sup>3</sup> | 1.727 ± 0.06 g/cm<sup>3</sup> |
| **ST** | 54.3 ± 3.0 dyne/cm | 63.5 ± 3.0 dyne/cm |
| **Polarizability** | 41.24 ± 0.5x10<sup>-24</sup> cm<sup>3</sup> | 20.99 ± 0.5x10<sup>-24</sup> cm<sup>3</sup> |
| **MR** | 104.04 ± 0.3 cm<sup>3</sup> | 52.97 ± 0.3 cm<sup>3</sup> |
| **MV** | 287.8 ± 3.0 cm<sup>3</sup> | 129.7 ± 3.0 cm<sup>3</sup> |
| **Nrotb** | 8 | 1 |
| **Drug likeness** | -3.95 | -6.29 |
| **Lipinski's rules Violations** | 0 | 0 |
| **SMILES** | COc2cc(C=CC(=O)CC(=O)C=Cc1ccc(O)c(OC)c1)ccc2O (E/Z)-Curcumin | [H]C(=O)c1c[nH]c2ccc(Br)cc12 |

MF= Molecular formula, MW= Molecular weight (Da), HBA= Hydrogen bond acceptors, HBD: Hydrogen bond donors, PSA= Polar Surface Area (Å<sup>2</sup>), LogP= Partition coefficient, δ= density (g/cm<sup>3</sup>), ST= Surface tension, MR= Molar refractivity (cm<sup>3</sup>), nrotb= No. of rotatable bonds, MV= Molar volume (cm<sup>3</sup>), SMILES= Simplified Molecular Input Line Entry System.

Molinspiration and Osiris Property Explorer simulators were used to predict the bioactivity of the compounds against important drug target sites and to predict their potential tumorigenic, irritant, reproductive and mutagenic risks, these results are found in Table 5. The bioactivity and biophysical properties indicate that curcumin is the compound with the greatest pharmacological interest, mainly at the target sites of nuclear receptor ligands and as an enzyme inhibitor, it is important that the values have a positive value to consider them as bioactive at a target site.

**Table 5.** Prediction of bioactivity and toxicity of compounds 1 and 2.

| *Compound* | **NCL** | **IE** | **GPCR** | **KI** | **PI** | **RE** | **T** | **M** | **I** |
| --- | --- | --- | --- | --- | --- | --- | --- | --- | --- |
| **Curcumin** | 0.12 | 0.08 | -0.06 | -0.26 | -0.14 | No | No | No | No |
| **5-bromoindole-3-carboxaldehyde** | -1.22 | -0.53 | -0.77 | -0.57 | -1.53 | No | No | No | No |

\* NCL= Nuclear receptor ligand, IE= Enzyme inhibitor, GPCR= GPCR ligand, KI= Kinase inhibitor, RE= Reproductive effective, T= Tumorigenic, M= Mutagenic, I= Irritant.

The greatest failure in the drug discovery process is toxicity and according to the prediction’s curcumin and 5-bromoindole-3-carboxaldehyde should not present tumorigenic, irritant, reproductive or mutagenic effects. The inhibition predictions of curcumin and 5-bromoindole-3-carboxaldehyde with CYP450 enzymes are made based on the structural formula of the compounds, this gives us an approximation of the metabolism of each molecule, according to table 5, curcumin inhibits the CYP3A4 and CYP1A2 isoforms, while carboxaldehyde only seems to inhibit CYP1A2, it is important to mention that the greater or lesser effect of an enzyme inhibitor will depend on factors such as the dose of the inhibitor and its capacity to bind to the enzyme, in the predictions made, it is not indicated whether the compounds are mild or potent inhibitors, but this data should be taken into account to consider possible drug–drug interactions (DDIs), as well as the doses that could be administered to a patient [47].

Based on the combined results of the cheminformatics properties, curcumin is the most promising compound to have the desired biological activity, which would be reflected in a possible better availability and distribution of the compound in the body, this is corroborated by the absorption data in Table 6.

**Table 6.** Data of ADME processes.

| *ADMET* |  | Curcumin | *5-bromoindole-3-carboxaldehyde* |
| --- | --- | --- | --- |
| **Absorption** | *WS* | -4.048 | -2.891 |
|  | *IA (%)* | 87.577 | 92.361 |
|  | *SP* | -2.755 | -2.425 |
|  | *SGP* | Yes | Yes |
|  | *P-GP-1-In* | Yes | No |
| **Distribution** | *BBBP* | -0.511 | Yes |
|  | *CNSP* | -3.07 | -1.758 |
| **Metabolism** | *CYP3A4 inhibitor* | Yes | No |
|  | *CYP1A2 inhibitor* | Yes | Yes |
| **Excretion** | *TC* | 0.06 | 0.204 |

\*WS: water solubility (log mol/L), IA: intestinal absorption, SP: skin permeability (logKp), SGP: P-glycoprotein substrate, P-GP-1-In: P-glycoprotein I inhibitor, BBBP: blood brain barrier permeability (logBB), CNSP: CNS permeability (logPS), TC: total clearance (log mL/min/kg).

### Molecular docking studies

We performed molecular docking studies to understand the binding interactions of curcumin and 5-bromoindole-3-carboxaldehyde with the LasR, PQS, and RhlR proteins of the Quorum sensing of *Pseudomona aeruginosa*. The molecular docking protocols were validated with re-docking, coupling the co-crystallized ligands in the proteins obtained in the Protein Data Bank using 12 methods from the combination of search algorithms and scoring functions, which are described in the materials and methods section. Figure 2 shows the results of the re-docking of the LasR, PQS and RhlR proteins with the co-crystallized ligands, as well as the method that was selected to perform the molecular docking of the two compounds under study from an RMSD less than 2 Å, which indicates that these methods can predict the experimental binding mode in the Quorum sensing proteins.

**Figure 2.**
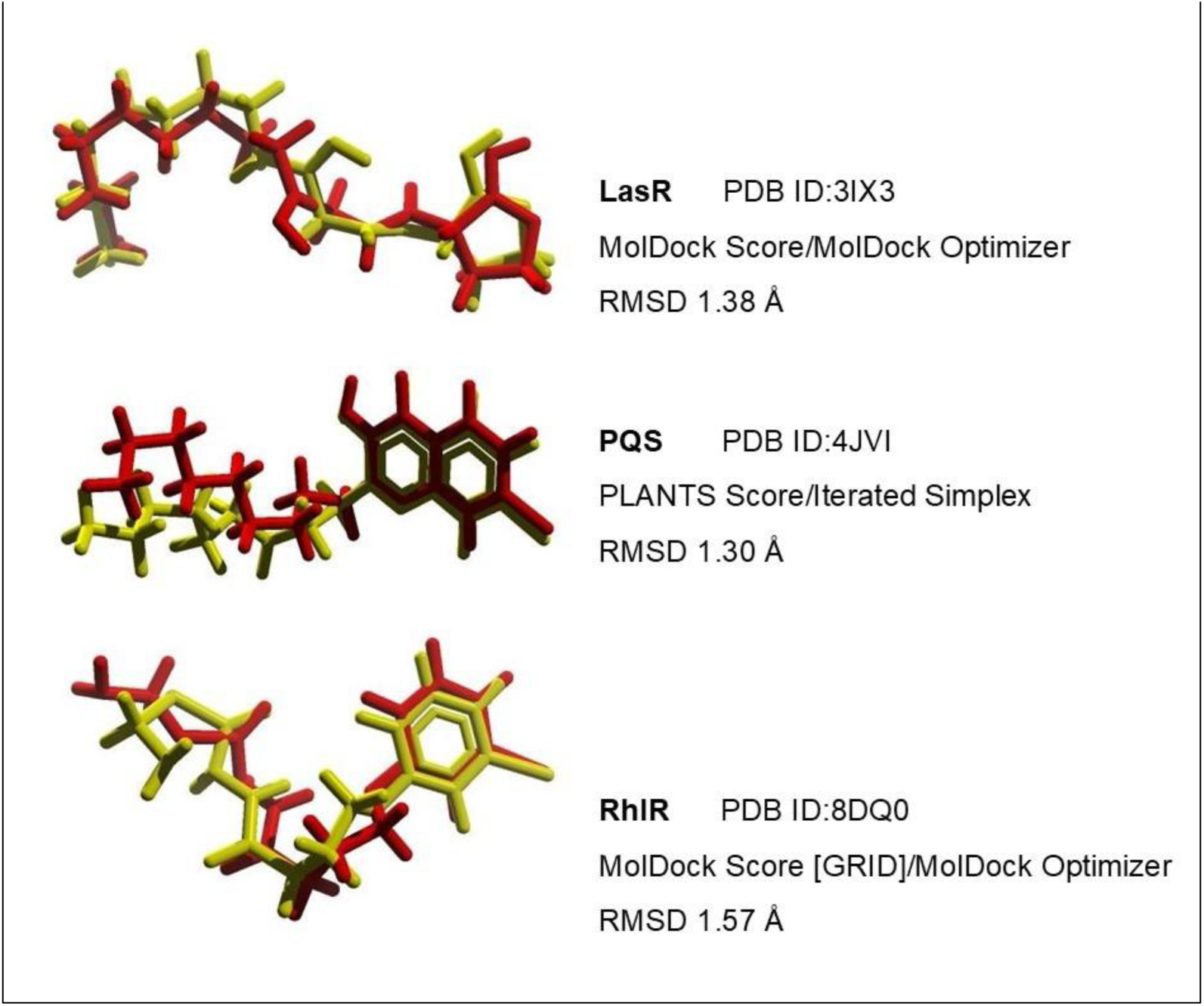
Re-docking analysis, the selected methods reproduce the pose of co-crystallized ligands (yellow colour) with RMSD less than 2 Å in the superposition of the re-coupled ligands (red colour). Docking protocols, each method is implemented by a scoring function and a search algorithm: LasR (MolDock Score/MolDock Optimizer), PQS (PLANTS Score/Iterated Simplex) and RhlR MolDock Score GRID/MolDock Optimizer).

Table 7 shows the affinity energies of curcumin and 5-bromoindole-3-carboxaldehyde on QS proteins, as well as the binding affinity of the endogenous inducers of the proteins LasR (3O-C12-HSL) and RhlR (C4-HSL) and the PQS inhibitor (3NH2-7Cl-C9-QZN) [48]. The amino acids of the ligand binding domain (LBD) with which the endogenous ligands interact are also shown, and the two compounds studied; as well as the types of ligand-receptor interaction. It should be noted that curcumin is the compound that has the highest affinity energy on the LBD in the 3 proteins, while 5-bromoindole-3-carboxaldehyde is the molecule with the lowest affinity energy; however, it is inferred that it could be a Quorum sensing inhibitor since it binds within the LBDs of the LasR, PQS and RhlR proteins, as can be seen in Figures 3, 4 and 5.

**Figure 3.**
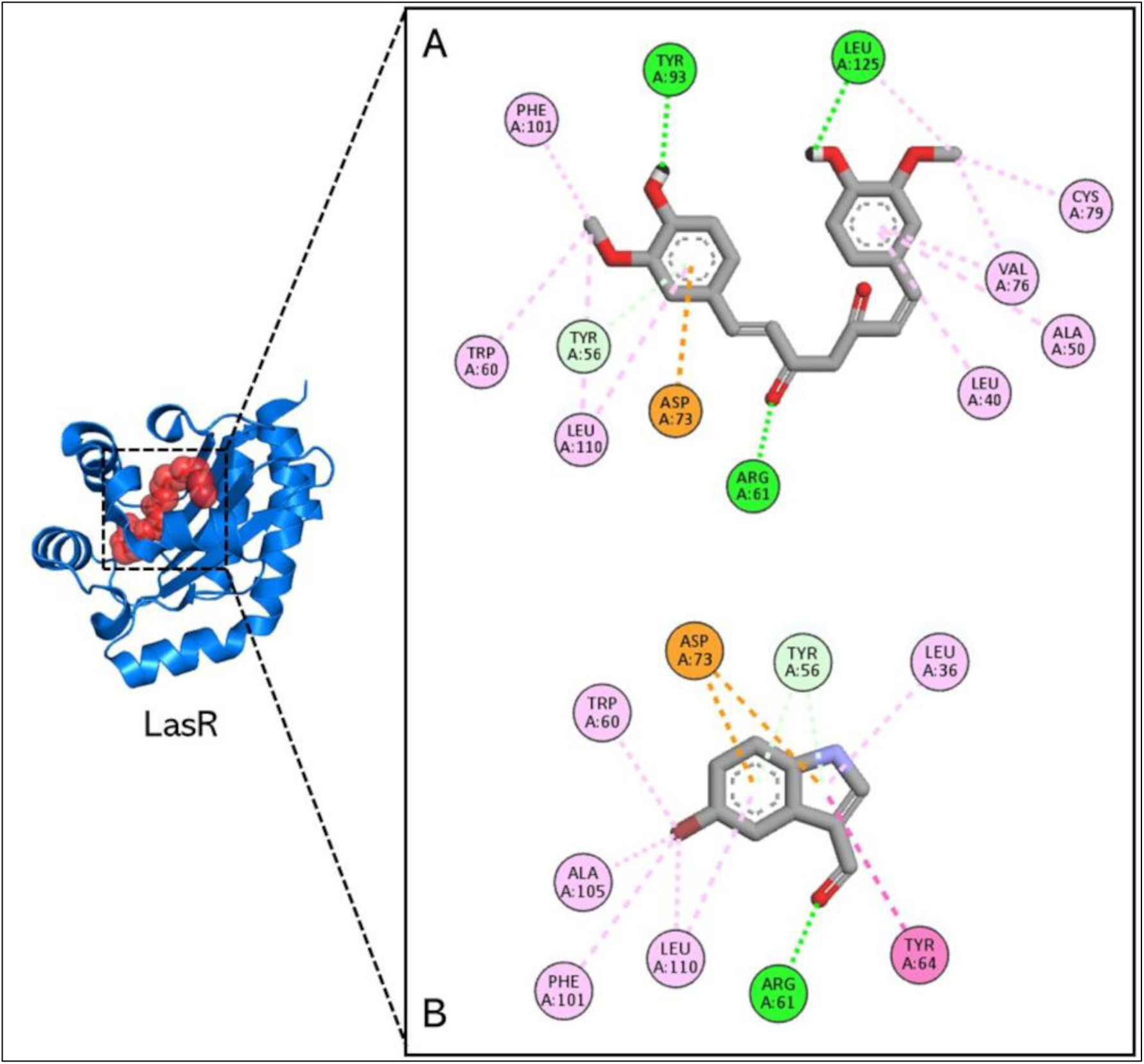
Results of molecular docking on LasR. The image shows the position of the ligand binding domain (red) on the LasR protein and the 2D interactions of curcumin (A) and 5-Bromoindole-3-carboxaldehyde (B).

**Figure 4.**
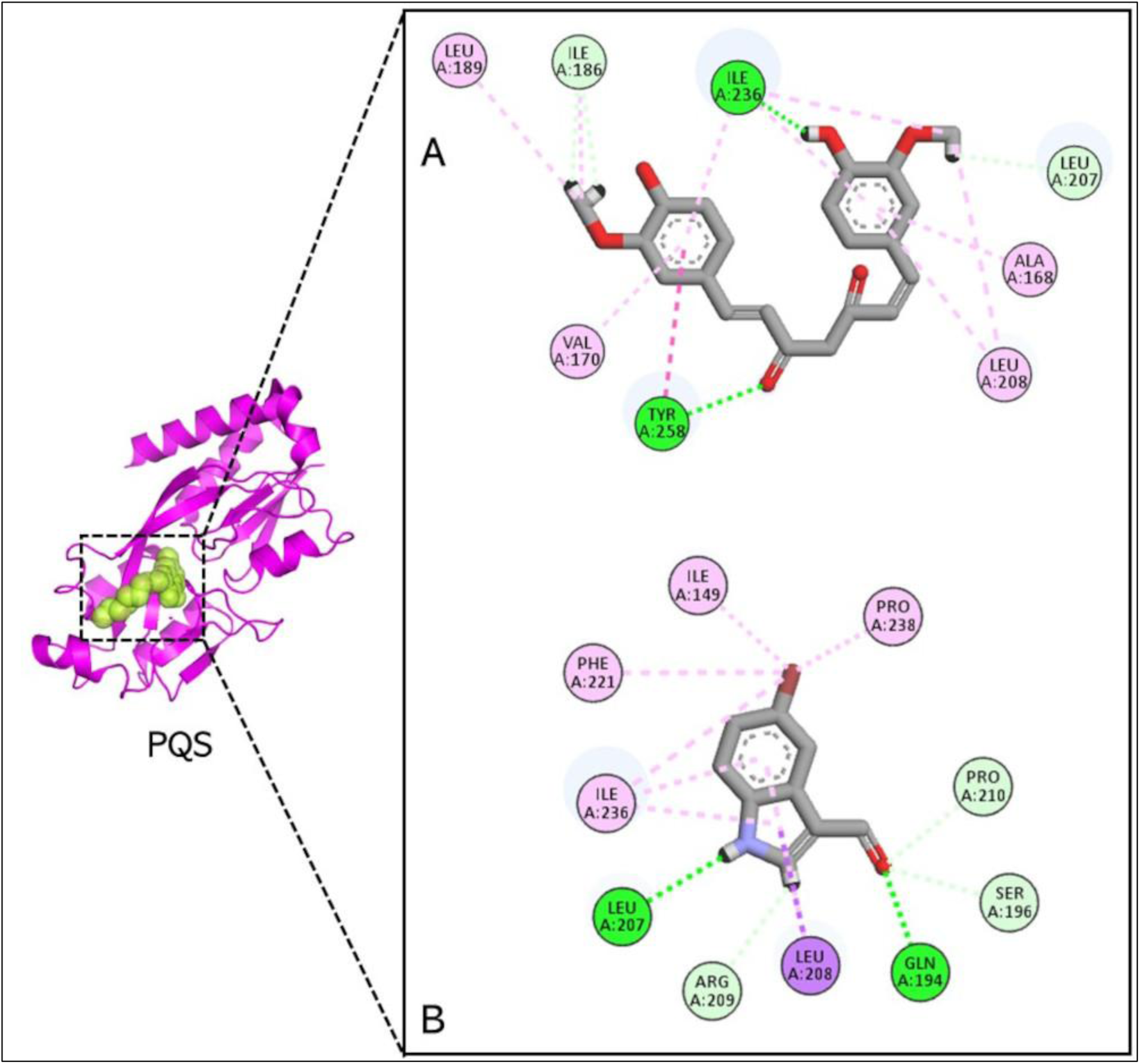
Results of molecular docking on PQS. The image shows the position of the ligand binding domain (yellow) on the PQS protein and the 2D interactions of curcumin (A) and 5-bromoindole-3-carboxaldehyde (B).

**Figure 5.**
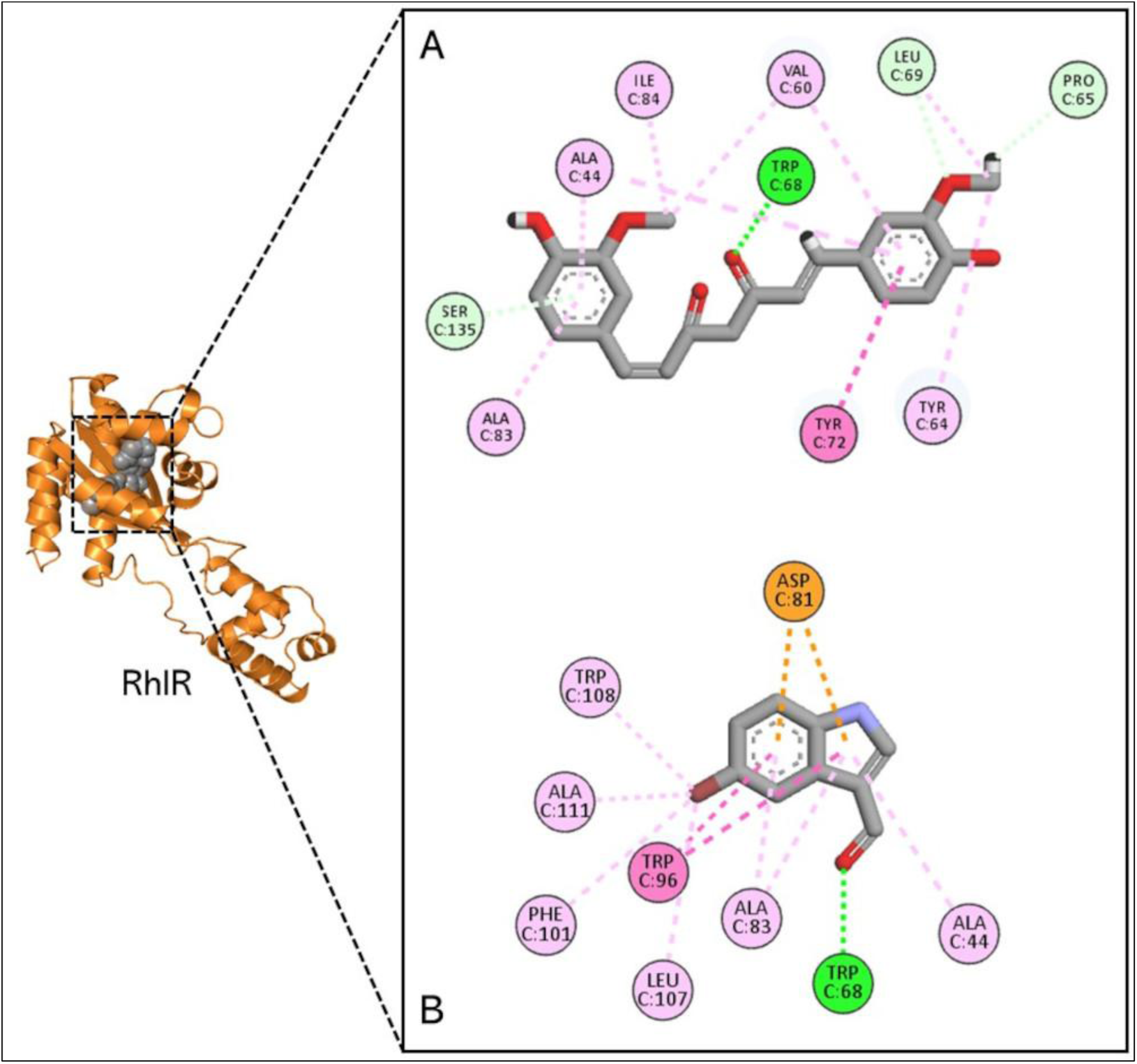
Results of molecular docking on RhlR. The image shows the position of the LBD (grey) on the RhlR protein and the 2D interactions of curcumin (A) and 5-bromoindole-3-carboxaldehyde (B).

**Table 7.** Affinity energy and binding mode of curcumin and 5-bromoindole-3-carboxaldehyde with *P. aeruginosa* Quorum sensing proteins.

| Target | Compounds | $\Delta G$<br>Kcal/mol | Interaction |
| --- | --- | --- | --- |
| LasR | 3O-C<sub>12</sub>-HSL | -9.29 | Hydrogen bond (Tyr56, Asp73, Thr75, Ser129), Carbon hydrogen bond (Ser29, Tyr93), alkyl (Leu36, Ala50, Val76, Cys79, Ala105, Leu110, Leu125, Ala127), pi-alkyl (Tyr64, Trp88, Phe101). |
|  | Curcumin | **-11.07** | Hydrogen bond (Arg61), pi-anion (Asp73), pi-donor hydrogen bond (Tyr56), pi-pi stacked (Tyr64), pi-pi T-shaped (Tyr56), alkyl (Ala105, Leu110), pi-alkyl (Leu36, Trp60, Phe101, Leu110). |
|  | 5-bromoindole-3-carboxaldehyde | -7.66 | Hydrogen bond (Arg61, Tyr93, Leu125), pi-anion (Asp73), pi-donor hydrogen bond (Tyr56), alkyl (Val76, Cys79, Leu125), pi-alkyl (Leu40, Ala50, Trp60, Val76, Phe101, Leu110). |
| PQS | 3NH<sub>2</sub>-7Cl-C9-QZN | -8.23 | Hydrogen bond (Leu207), alkyl (Ala102, Ile149, Ile186, Ile189, Leu207, Ile236, Pro238, Ile263), pi-alkyl (Ile149, Ala168, Leu207, Leu208, Ile236, Tyr258). |
|  | Curcumin | **-9.23** | Hydrogen bond (Gln194, Leu207), carbon hydrogen bond (Ser196, Arg209, Pro210), pi-sigma (Leu208), alkyl (Ile149, Ile236, Pro238), pi-alkyl (Leu208, Phe221, Ile236). |
|  | 5-Bromoindole-3-carboxaldehyde | -7.14 | Hydrogen bond (Ile236, Tyr258), carbon hydrogen bond (Ile186, Leu207), pi-pi T-shaped (Tyr258), alkyl (Ile186, Leu189, Leu208, Ile236), pi-alkyl (Ala168, Val170, Leu208, Ile236). |
| RhIR | C4-HSL | -7.12 | Hydrogen bond (Try64, Trp68, Asp81, Ser135), alkyl (Ile84, Leu107, Ala111, Val133), pi-alkyl (Trp96, Phe101). |
|  | Curcumin | **-10.27** | Hydrogen bond (Trp68), pi-anion (Asp81), pi-pi stacked (Trp96), alkyl (Leu107, Ala111), pi-alkyl (Ala44, Ala83, Phe101, Trp108). |
|  | 5-bromoindole-3-carboxaldehyde | -7.68 | Hydrogen bond (Trp68), carbon hydrogen bond (Pro65, Leu69), pi-donor hydrogen bond (Ser135), pi-pi T-shaped (Tyr72), alkyl (Val60, Leu69, Ile84), pi-alkyl (Ala44, Tyr64, Val60, Ala83). |

The ligand binding domain of LasR is housed in a large hydrophobic pocket formed by the amino acids Leu36, Gly38, Leu39, Leu40, Tyr47, Glu48, Ala50, Ile52, Tyr56, Trp60, Arg61, Tyr64, Asp65, Gly68, Tyr69, Ala70, Asp73, Pro74, Thr75, Val76, Cys79, Thr80, Trp88, Tyr93, Phe101, Phe102, Ala105, Leu110, Thr115, Leu125, Gly126, Ala127, and Ser129 [49]. Figure 3 shows the interactions between curcumin and 5-bromoindole-3-carboxaldehyde with LBD amino acids and that they share with the endogenous ligand 3O-C12-HSL, such as Tyr56, Asp73, Leu110 and Phe101. In addition, curcumin also interacts with Leu36 and Tyr64 as the endogenous ligand. Meanwhile, 5-bromoindole-3-carboxaldehyde as well as 3O-C12-HSL interact with the residues Ala50, Val76, Cys79, Tyr93 and Leu125.

Figure 4 shows the structure of the LysR-like transcription factor that responds to the quinolone signal in *Pseudomonas* (PQS) present in *Pseudomona aeruginosa*, as well as the ligand binding domain in yellow that covers from residue 91 to 319 [50]. Ilangovan et al., in 2013 evaluated quinazolinone analogues and reported the compound 3NH2-7Cl-C9-QZN (QZN 23) as a novel PQS inhibitor with an IC50 of 5.0 μM [48], which binds to the LBD with the amino acids reported in Table 7. The interactions between curcumin and 5-bromoindole-3-carboxaldehyde in the LBD are shown in Figure 4. QZN-23 inhibitor, curcumin, and 5-bromoindole-3-carboxaldehyde interact with Leu207, Leu208, and Ile 236 residues. Curcumin also interacts with the amino acids Ile149 and Pro238 as well as the inhibitor QZN 23. While 5-bromoindole-3-carboxaldehyde like QZN 23 binds with Ala168, Ile186, Ile189 and Tyr258.

It has been reported that for the multimerization and activity of RhlR, its autoinducer C4-HSL is necessary, which binds in the LBD efficiently and that, in its absence, homodimerizes decreasing its capacity as a transcriptional activator [51]. The residues of RhlR LBD with which C4-HSL interacts are reported in Table 7. Both C4-HSL, curcumin and 5-bromoindole-3-carboxaldehyde bind in LBD and interact commonly with Trp68. Curcumin also binds with other amino acids with which it interacts with C4-HSL, Asp81, Phe101, Leu107 and Ala111. Meanwhile, 5-bromoindole-3-carboxaldehyde binds with Tyr64, Ile84 and Ser135 residues as well as C4-HSL, as shown in Figure 5.

### Molecular dynamics simulations

Since molecular docking does not account the interactions with the environment surrounding the complexes, the thermodynamic effects, or the intrinsic flexibility of the protein and ligand, we conducted 200-ns MD simulations to further assess the dynamic stability and conformational behavior of the protein–ligand systems. Analyses of hydrogen bonds, root RMSD, RMSF and an appropriate estimate of the binding energy using the MM/PBSA method, allowed us a more comprehensive of the interactions and validating predictions obtained from molecular docking for complexes involving curcumin and 5-bromoindole-3-carboxaldehyde.

Hydrogen bonds are critical for maintaining the stability of protein-ligand complexes. Figure 6 shows the number of hydrogen bonds formed between curcumin or 5-bromoindole-3-carboxaldehyde and each of the three quorum-sensing proteins over 200 ns of simulation time in MD. The complexes in which curcumin is involved (Figures 6 A-C) display a relatively higher and more fluctuating number of hydrogen bonds across the simulation. In particular, the LasR–curcumin suggests a dynamic but consistently engaged interaction network (Figure 6 A). The PQS-related protein complex (Figure 6 B) shows a gradual increase in hydrogen-bond formation after approximately 120 ns, which may indicate a progressive stabilization or rearrangement of the ligand within the binding pocket. In contrast, the RhIR–curcumin complex (Figure 6 C) exhibits more pronounced fluctuations and intermittent hydrogen bonding, advising a less stable or more transient interaction pattern. For 5-bromoindole-3-carboxaldehyde (Figures 6 D-F), the number of hydrogen bonds is generally lower. Notably, the RhIR–5-bromoindole-3-carboxaldehyde complex (Figure 6 F) shows the most persistent hydrogen-bonding behavior through simulation time, suggesting a relatively stable binding mode in this specific protein.

**Figure 6.**
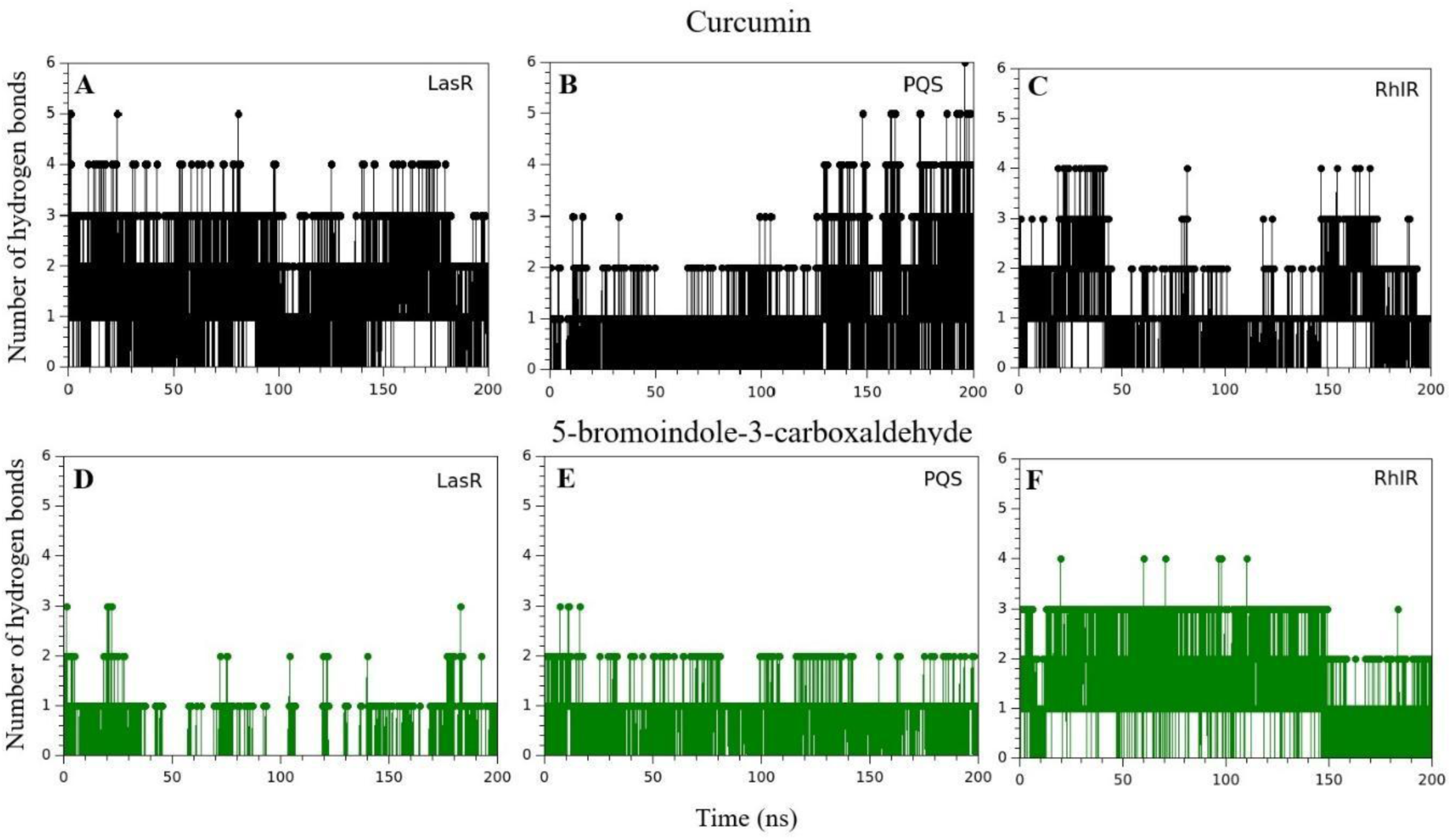
Number of hydrogen bonds as a function of time over 200 ns for the 6 different complexes: **A** LasR-curcumin; **B** PQS-curcumin; **C** RhIR-curcumin; **D** LasR-5-bromoindole-3-carboxaldehyde; **E** PQS-5-bromoindole-3-carboxaldehyde; and **F** RhIR-5-bromoindole-3-carboxaldehyde.

RMSD was also obtained from MD simulations. RMSD provides information on the average atomic displacement between two different conformations by least-square fitting the structure over simulation time to the reference structure, so it is commonly used to assess the structural stability of protein–ligand complexes. Figure 7 presents the RMDS over 200 ns of MD simulation for the protein backbones and the compounds (curcumin and 5-bromoindole-3-carboxaldehyde) in the complexes. For the backbone of each protein, the RMSD in Figure 7 A indicates that LasR and PQS exhibit stable conformations from early times. In contrast, RhIR shows slight fluctuations for the complex with curcumin, stabilizing after ∼100 ns. However, for the complex with 5-bromoindole-3-carboxaldehyde, RhIR displays pronounced RMSD fluctuations, reaching ∼1.2–1.4 nm during the simulation. This RMSD profile suggests enhanced conformational flexibility or partial rearrangement of the protein structure, which tends to stabilize at longer times, indicating that the systems eventually reach dynamic equilibrium. In Figure 7 B, the heavy-atom RMSD of curcumin with respect to protein backbone shows a structurally stable conformation for this compound within each of the three proteins throughout the entire simulation time. On the other hand, the heavy-atom RMSD of 5-bromoindole-3-carboxaldehyde with respect to protein backbone shows markedly different behavior depending on the protein. While the ligand remains relatively stable in the PQS and RhlR binding sites, its RMSD in the LasR complex increases significantly, reaching values above ∼6 nm. This large deviation suggests substantial 5-bromoindole-3-carboxaldehyde mobility, possible reorientation, or partial dissociation from the binding pocket, indicating weaker or less stable binding to LasR.

**Figure 7.**
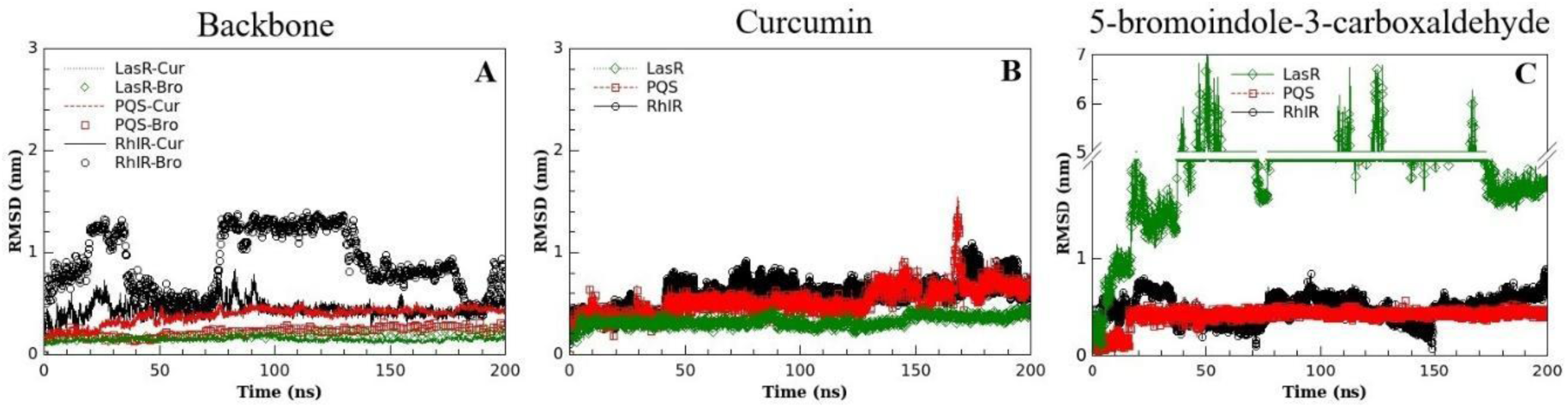
Root mean square deviation (RMSD) over the 200 ns of MD simulations: **A** Analysis of protein backbone respect to the backbone itself (lines or open symbols indicate the presence of curcumin or 5-bromoindole-3-carboxaldehyde, respectively); **B** Analysis of heavy-atoms of curcumin respect to protein backbone; and **C** Analysis of heavy-atoms of 5-bromoindole-3-carboxaldehyde respect to protein backbone. In all cases, green is for LasR, red for PQS, and black for RhIR.

RMSF complements the structural stability analysis of complexes. RMSF is time-independent but rather the fluctuation of the displacement of an atom relative to its average position over time. For curcumin (left panel of Figure 8), the RMSF analysis shows that, although atomic fluctuations below ∼0.15 nm indicate a certain degree of flexibility, most regions remain nearly rigid across all proteins, suggesting that curcumin is relatively well anchored within the binding sites. Nevertheless, distinct regions of enhanced flexibility are observed, particularly at the terminal aromatic rings. These segments display higher RMSF values, most notably in the RhIR complex. This behavior implies that while the central scaffold of curcumin is stably bound, its terminal groups undergo conformational adjustments that may enable adaptive interactions with the diverse quorum-sensing protein environments. For 5-bromoindole-3-carboxaldehyde, on the right panel of Figure 8, it can be observed that it exhibits substantially lower RMSF across all atoms and protein complexes. Most atoms fluctuate below ∼0.05 nm, indicating a rigid binding mode and limited internal flexibility. A slight increase in RMSF is observed for atoms located close to the aldehyde functional group, particularly in the LasR and PQS complexes, which may reflect localized adjustments to optimize hydrogen bonding or electrostatic interactions.

**Figure 8.**
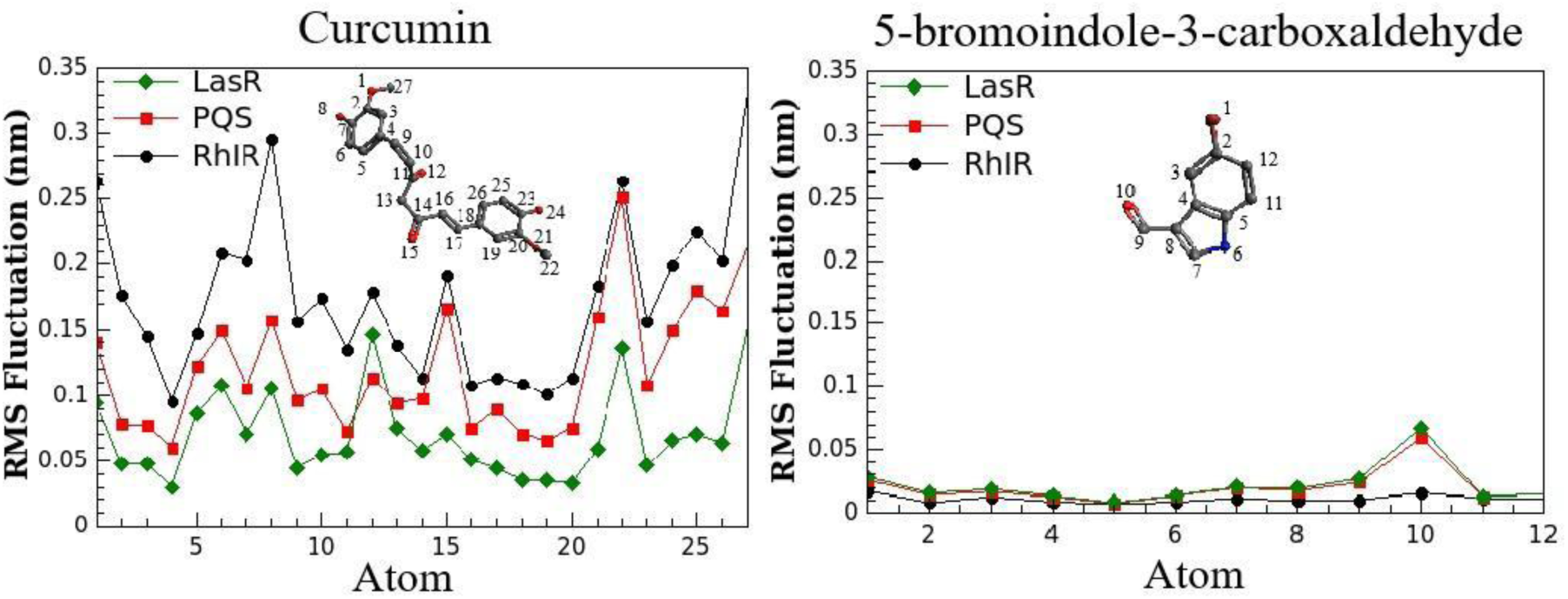
Root mean square fluctuation over 200 ns for the atoms of curcumin (left panel) and 5-bromoindole-3-carboxaldehyde (right panel) when the ligands form complexes with the proteins LasR (green diamonds), PQS (red squares) or RhIR (black circles). The atom number correspondence is according to the structure presented inside each figure.

Figure 9 shows RMSF of LasR, PQS and RhIR in presence of curcumin or 5-bromoindole-3-carboxaldehyde. For LasR (Figure 9 A), both complexes show relatively low RMSF, which indicates an overall rigid and stable protein structure during the simulation. In PQS (Figure 9 B) moderate fluctuations are observed in several regions, being more evident for curcumin. However, the RMSF remains stable and no ligand compromises the overall stability of the PQS protein. In the case of RhIR, Figure 9 C shows pronounced differences for the two ligands. The complex with 5-bromoindole-3-carboxaldehyde exhibits significantly higher RMSF values, particularly in the C-terminal region, where fluctuations exceed 1 nm. This indicates substantial conformational flexibility of the protein.

**Figure 9.**
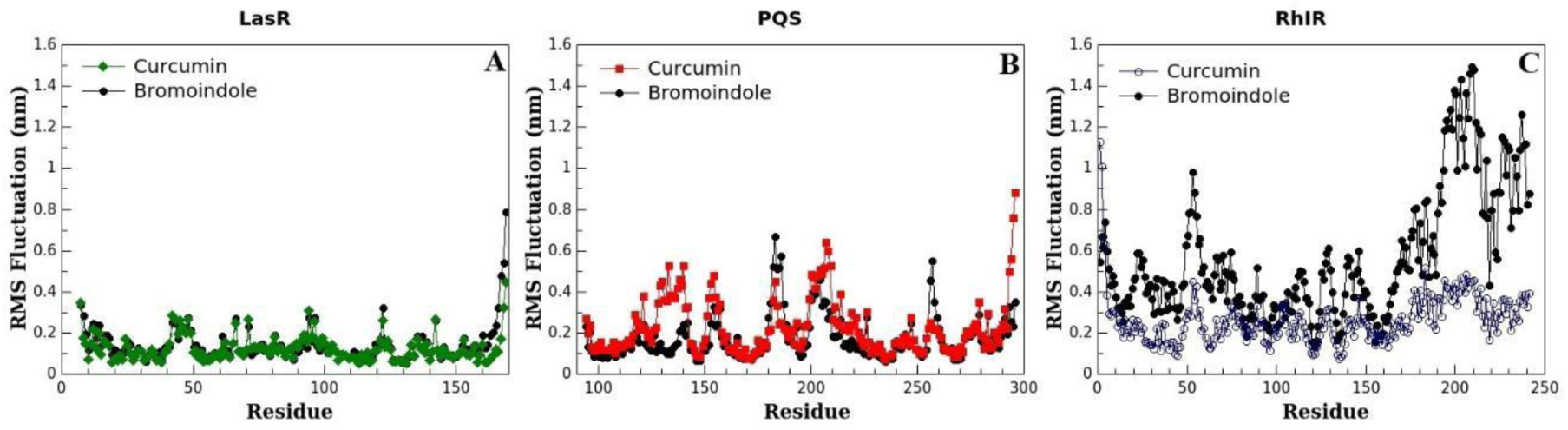
Root mean square fluctuation over 200 ns of the aminoacids in proteins: **A** LasR in presence of curcumin (green diamonds) or 5-bromoindole-3-carboxaldehyde (black circles); **B** PQS in presence of curcumin (red squares) or 5-bromoindole-3-carboxaldehyde (black circles); and **C** RhIR in presence of curcumin (blue open circles) or 5-bromoindole-3-carboxaldehyde (black circles).

To quantify the strength of the interaction in complexes, we extend our study by providing a more accurate estimate of the binding energy by means of the MMPBSA approximation. The Gibbs free energy aims to encompass the results obtained by considering the thermodynamic stability of the complexes. In this manner, Table 8 shows electrostatic, van der Waals, polar and non-polar contributions to the total binding free energy. As can be seen, in all complexes, gas phase free energy dominates the binding. Of these, van der Waals interactions are predominant and responsible for the stability of the compounds in the binding pockets due to the presence of hydrophobic contacts. Although electrostatic interactions are also attractive, their contribution is not as significant, so the stability of the complexes is not dictated entirely by the formation of hydrogen bonds. For all systems, polar interactions significantly reduce the affinity between proteins and ligands, reflecting the energy penalty associated with loss of favorable protein/ligand-solvent interactions upon binding. Overall, the binding energy calculations indicate that all protein-ligand complexes are thermodynamically favorable and stable. However, complexes with 5-bromoindole-3-carboxaldehyde exhibit weaker van der Waals and electrostatic interactions, except in the case of RhIR where the electrostatic contribution is more favorable than that observed for curcumin. Although the polar contribution is lower than curcumin, reflecting reduced desolvation of polar groups, this effect does not compensate for the significantly weaker intermolecular interactions. Notably, the results demonstrate that curcumin presents stronger and more stable binding, validating it as the compound with the highest affinity in the three *P. aeruginosa* quorum-sensing proteins.

**Table 8.** MM/PBSA binding free energy results expressed in units of kcal/mol.

| Target | Complex<br>Compound | Binding free energy terms |  |  |  |  |
| --- | --- | --- | --- | --- | --- | --- |
| | | $\Delta E_{vdw}$ | $\Delta E_{ele}$ | $\Delta E_{pol}$ | $\Delta E_{non-pol}$ | $\Delta G_{bind}$ |
| LasR | Curcumin | -49.43±2.91 | -14.90± 9.21 | 49.33±8.93 | -4.86±0.14 | **-19.86±6.38** |
|  | 5-bromoindole-3-carboxaldehyde | -10.13±7.05 | -4.75±7.80 | 8.75±10.42 | -1.55±0.88 | **-7.67±5.46** |
| PQS | Curcumin | -37.88±4.16 | -14.02±6.05 | 31.12±6.21 | -4.62±0.25 | **-25.39±6.01** |
|  | 5-bromoindole-3-carboxaldehyde | -20.00±1.84 | -5.89±4.19 | 17.12±4.31 | -2.48±0.10 | **-11.26±5.42** |
| RhIR | Curcumin | -41.33±4.74 | -12.01±5.91 | 33.87±6.62 | -4.58±0.22 | **-24.05±6.13** |
|  | 5-bromoindole-3-carboxaldehyde | -21.08±2.67 | -16.18±6.32 | 28.46±3.58 | -2.47±0.07 | **-11.26±6.10** |

## Discussion

The minimum inhibitory concentrations obtained for curcumin correspond to those reported by Adamczak [21] for Pa01 and ATCC 27853, and Gunes [52] for strain ATCC 27853, as well as those described by Ramesh [53] for Pa01 and ATCC 27853, and Gholami [54] for Pa01. The concentrations in previous studies ranged from 125 to 2000 µg/mL. This shows that the concentrations evaluated for quorum inhibition correspond to sub-inhibitory concentrations and are comparable to those already reported.

On the other hand, for 5-bromoindole-3-carboxaldehyde, there are no reported MIC studies in Pseudomonas aeruginosa. Therefore, we started with the highest dose evaluated for quorum inhibition in our work, up to 1000 µg/mL, which also demonstrates that we worked with sub-inhibitory concentrations. The concentrations of 30, 50, and 1000 µg/mL were chosen because the same carboxaldehyde was studied in *C. violaceum* in a study by Kemp [18] to evaluate quorum inhibition.

It has been shown that *P. aeruginosa* employs quorum sensing in the regulation of genes encoding extracellular virulence factors. Quorum sensing is an intercellular communication system that allows bacteria to control gene expression in a cell population density-dependent manner [55].

In *P. aeruginosa*, four fundamental QS pathways have been identified. The first is known as the LasI/LasR cascade. The second includes the RhlI/RhlR pathway. The third is the *Pseudomonas* quinolone signal (PQS). The fourth is the AI-based integrated quorum sensing system [56].

In particular, QS blockade abrogates the virulence without affecting bacterial growth. Therefore, QS represents an attractive target for anti-virulence therapy [57].

To contextualize our findings with what has been reported in the literature, Table 9 summarizes previous studies on curcumin and bromoindole, both in terms of MIC and quorum sensing inhibition. This comparison highlights the consistency of our experimental design and the main differences with established studies, and underscores the novelty of the MIC assessment for 5-bromoindole-3-carboxaldehyde in *P. aeruginosa*.

**Table 9.**
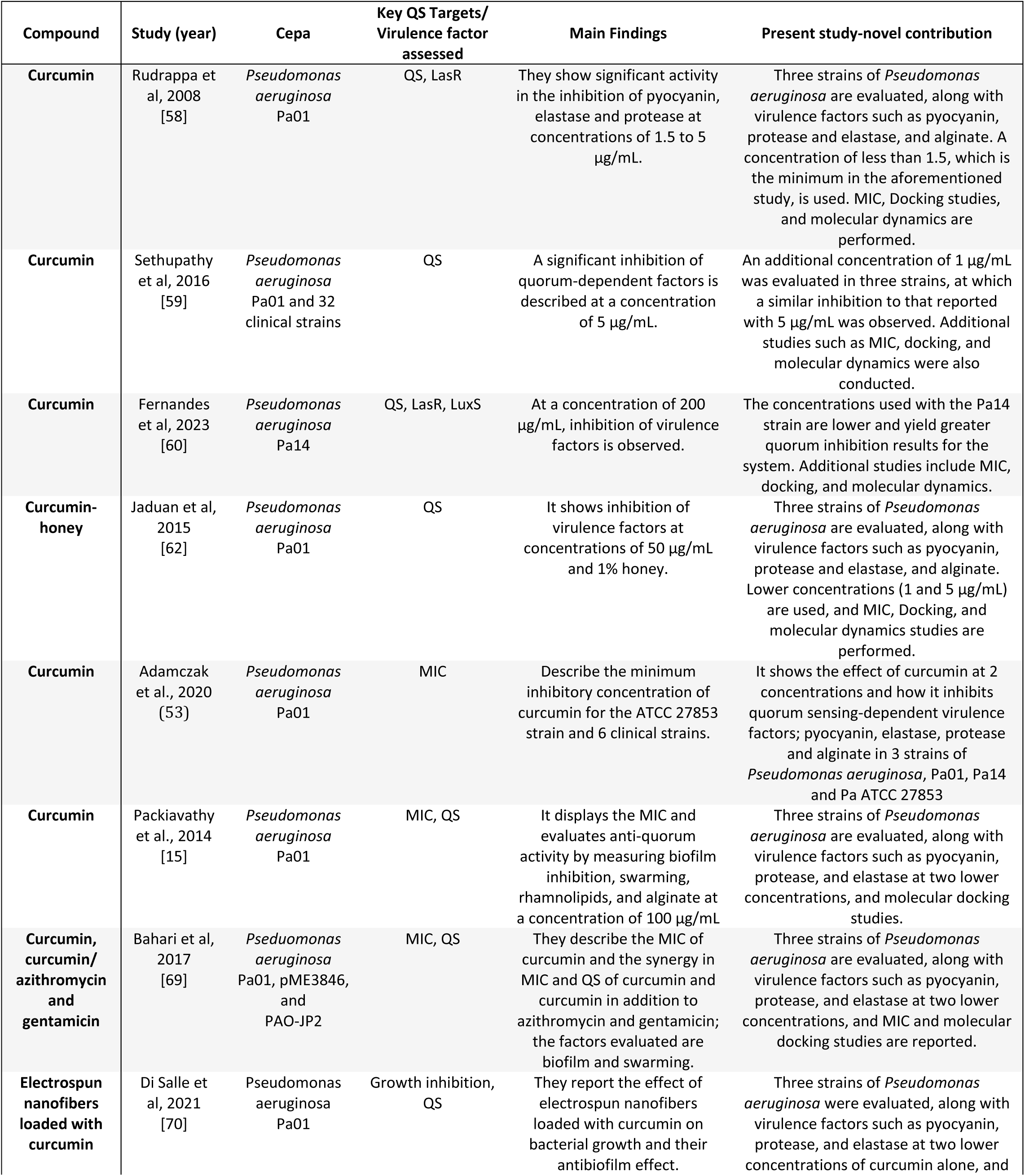
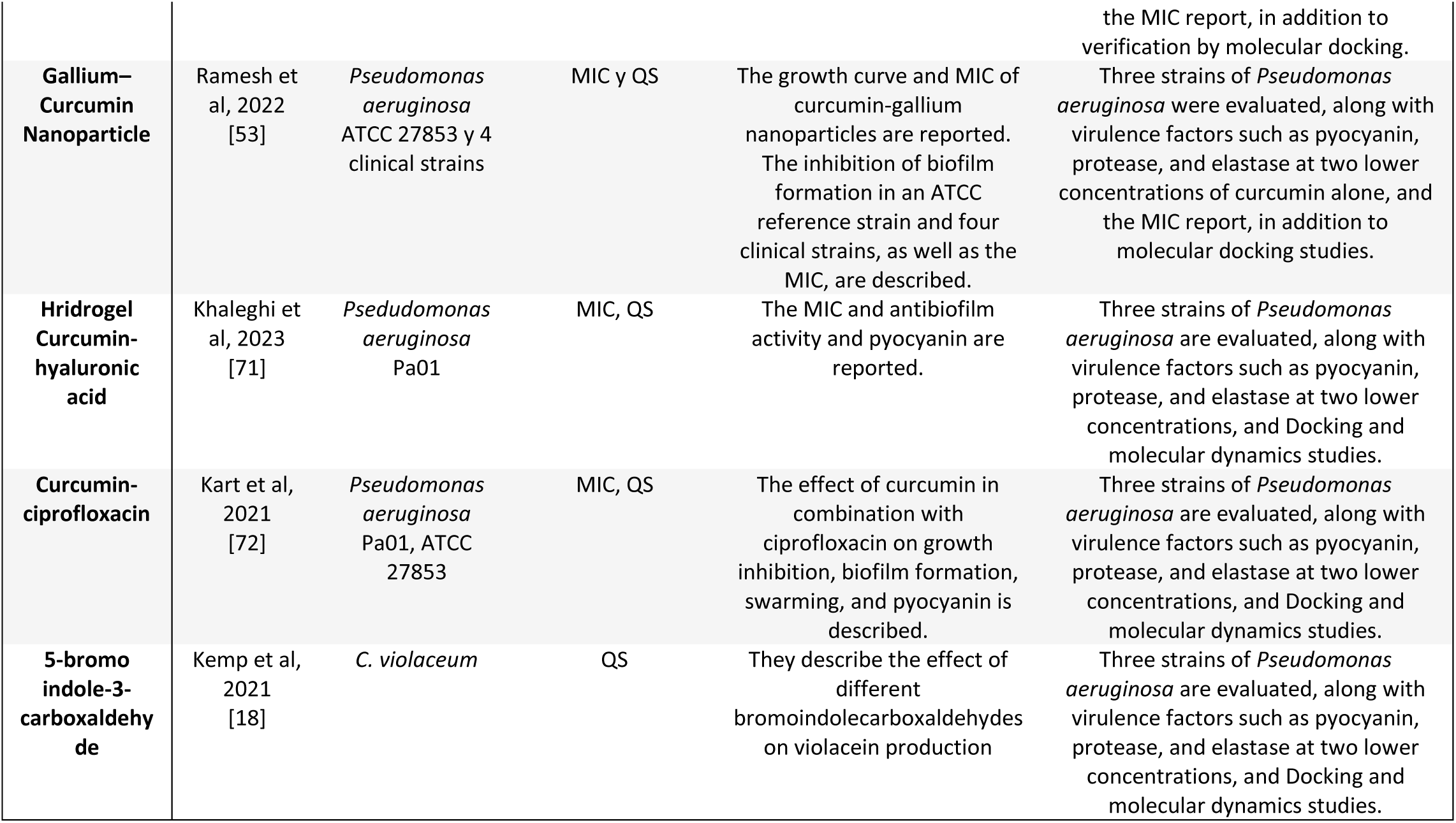
Reported MIC and quorum-sensing inhibition in *Pseudomonas aeruginosa*.

In a study conducted by Rudrappa in Pa01 where they evaluated different concentrations of curcumin, at concentrations of 1.5 to 3 µg/ml they achieved an inhibition of pyocyanin of between 60 and 80% of the production with respect to the control, and at 5 µg/ml they achieved an inhibition of 3 times compared to the control. Not only did they achieve an inhibition of pyocyanin, other factors regulated by the QS systems lasI-lasR and rhlI-rhlR were also studied, where they observed a double decrease in the production of elastase and protease, where they also observed that curcumin directly affected the production of the lactones 3-oxoC12-HSL and C4-HSL. In our study we observed that at 1 µg/ml of curcumin we obtained an inhibition for Pa01 of 60% which is similar to that observed for Rudrappa, a value similar to that observed in the ATCC 27853 strain [58].

Sethupathy and collaborators demonstrated the activity of curcumin at 5 µg/ml on the inhibition of pyocyanin, elastase, protease, alginate and biofilm in 32 clinical isolates and using Pa01 as a control, finding inhibitions that they considered as effective inhibition (50-90%), minimal inhibition (20-49%), no inhibition (0-19%) and induced. Where most of the strains had an effectiveness of between 50 and 90% at said concentration, the factors that showed said inhibition are mainly pyocyanin and elastase. The rest of the strains were maintained with minimal inhibition [59]. In our study at a concentration of 1 µg/ml for Pa01, an inhibition of 60.60% was obtained and for 5 mg of 90.63%, for Pa14 an inhibition of 55.89% was achieved at that concentration, where the greatest activity is seen is in the ATCC 27853 strain where it is achieves 100% pyocyanin inhibition. In addition to similar results in the rest of the virulence factors.

A published study reported for Fernandes, that curcumin has inhibitory effects on the LuxS/AI-2 and LasI/LasR QS systems, resulting in a reduction of 33%–77% and 21%, respectively. Significantly, curcumin at high concentration of 200 μg/ml resulted in a 21% decrease in the generation of 3-oxo-C12-HSL by *P. aeruginosa* Pa14 strain [60].

Liu and collaborators, they monitored pyocyanin production and biofilm formation in *Pseudomonas aeruginosa* when adding curcumin, and they showed a significant decrease in both virulence factors with the molecule [61].

Gholami and collaborators studied the activity of curcumin and curcumin with metal complexes to inhibit quorum and found important decreases in pyocyanin and considerable alginate. Their study shows that curcumin alone reduces pyocyanin production by more than half with a 67.5 % compared to the control in Pa01, and to a lesser extent with metals. On the other hand, the activity in alginate with curcumin alone is lower than that shown in combination with metals with only 21.3% [54].

Jaduan tested the effect of curcumin mixed with honey to inhibit quorum in *Pseudomonas aeruginosa* in strain Pa01 with the following results: Treatment with curcumin (50 µg/ml) and honey (1%) only showed a weak inhibitory effect on siderophore production, a result similar to that observed in treatment with curcumin alone. Curcumin (50 µg/ml) and honey (1%) alone were not found to significantly reduce the activity of Pa01 metalloproteases elastase y protease. Interestingly, curcumin (50 µg/ml) and honey (1%) alone were ineffective in decreasing alginate production [62]. This synergy shows lower activity than what we obtained in our study for the virulence factors evaluated.

It is known that curcumin acts on the las system. This signalling system is regulated by virulence factors such as the expression of alkaline protease and elastase, which represents the decrease in the production of the autoinducer lactone and therefore the inhibition of production of said factors. As shown in the results [65]. If lasR/lasI system transcriptional regulators expression is affected, other systems such as the PQS and all virulence factors in *P. aeruginosa* will be affected too [63].

Several studies have described the antibiofilm and antivirulence activities of indole derivatives against clinically important pathogens such as *Pseudomonas aeruginosa* [65]. Halogenation of indoles demonstrates a decrease in IC50 and also greater potency as inhibitors of quorum sensing, as described by Odularu et. al., [17]. The antimicrobial activity of some indolecarboxaldehydes has been demonstrated in strains of *Pseudomonas sp*, but there is no information on quorum inhibition in *Pseudomonas aeruginosa* [66].

Among the most studied bacterial species, *C. violaceum* has been shown to affect the production of AHL with good results at concentrations similar to those studied by us [18].

Indolecarboxaldehydes such as 5-bromoindole-3-carboxaldehyde and derivatives in *Vibrio fischeri* have also been shown to inhibit the LuxI/LuxR system, thereby quenching its bioluminescence or biofilm inhibition in *Vibrio parahaemolyticus* [18,67].

In *Pseudomonas aeruginosa*, the following indole derivatives have been found Indole, 3-indolylacetonitrile, 7-fluoroindole, indole-based N-acylated L-homoserine lactones and 2-Aminobenzimidazoles, which can inhibit the production of virulence factors and biofilm formation, however, carboxaldehyde derivatives with this activity are not reported [68].

The solvent did not affect the production of virulence factors at concentrations of 0.01%, 0.5% and 1% (v/v), according to the prepared stocks of the molecules, since equal results were obtained in control and negative control, however some authors report that DMSO can act on quorum sensing at higher concentrations such as 2% (v/v).

The analyses of variance performed for elastase, protease and alginate activities in *Pseudomonas aeruginosa* with Curcumin and 5-bromoindole-3-carboxaldehyde revealed significant differences only in protease activity between treatments. These results suggest that while some compounds may influence the activity of certain enzymes, not all treatments have a uniform impact on all enzyme activities evaluated.

Some limitations of our work are molecular studies to know exactly which gene is repressed and to what extent it is done, however, this represents an opportunity for the future, especially in the case of bromoindole studied, since so far, we are the first to report this activity in three different strains of *Pseudomonas aeruginosa*.

Chemoinformatic properties related to ADME processes were predicted using the online predictors pkCSM and SwissADME (Table 6). Absorption properties indicate that the compounds can be administered orally according to Lipinsky rules. Other predicted properties, such as LogP data, absorption, highlighting intestinal absorption which is 87.5% for curcumin and 92.3% for 5-BIC, the skin permeability of both compounds is relatively similar (−2.755 and −2.425 respectively), in addition to the fact that curcumin could inhibit P-GP-1-In, which would cause the compound not to be expelled from the cell and could promote biological activity. On the other hand, 5-BIC appears to permeate the blood-brain barrier while curcumin does not, this is to be expected due to the difference in size, polar atoms and solubility, but opens the possibility that 5-BIC could be used in the treatment of infections affecting the Central Nervous System.

*In silico* analysis was used to support the hypothesis that curcumin and 5-bromoindole-3-carboxaldehyde interact with the three major Quorum sensing proteins of *P. aeruginosa* (Table 7 and Figures 3-5). Molecular docking studies were performed to obtain the affinity energy and binding mode of the molecules under study in the LBDs of the pharmacological targets.

Molecular docking with the LasR protein showed that curcumin has a stronger binding (−11.07 Kcal/mol) compared to its endogenous ligand (−9.29 Kcal/mol), while 5-bromoindole-3-carboxaldehyde was weaker in affinity (−7.66 Kcal/mol). Similarly, curcumin has a higher energy affinity for the PQS protein (−9.23 Kcal/mol) compared to the QZN 23 ligand that has an IC_50_ of 5.0 μM (−8.23 Kcal/mol), which we rely on to infer that curcumin could be a PQS inhibitor, while 5-bromoindole-3-carboxaldehyde has a weaker binding (−7.14 Kcal/mol). Finally, with the RhlR protein, curcumin also presented a higher affinity energy (−10.27 Kcal/mol) compared to the affinity presented by the endogenous ligand C4-HSL (−7.12 Kcal/mol), while 5-bromoindole-3-carboxaldehyde presented an affinity like the endogenous ligand (−7.68 Kcal/mol). It is important to mention that the result of the affinity energy is only indicative of a higher probability of a more stable bond. Therefore, the binding mode of the ligands under study should be compared with endogenous ligands or other inhibitors or antagonists of the Quorum sensing proteins.

In this sense, Figures 3, 4 and 5 show the binding modes of curcumin and 5-bromoindole-3-carboxaldehyde in the ligand binding domains of the LasR, PQS and RhlR proteins. It is shown that both compounds under study can accommodate at the site of activation of LasR and RhlR, suggesting a possible competition or antagonism with their autoinducers in the LBD and by interacting with essential residues they promote a disruption of the QS signalling cascade. Likewise, curcumin and 5-bromoindole-3-carboxaldehyde are accommodated in the same LBD cavity as the QZN 23 inhibitor and bind with amino acids relevant to PQS activity, preventing their interaction with DNA and impeding their transcriptional activity. Taken together, the *in silico* analysis supports experimental results showing that both curcumin and 5-bromoindole-3-carboxdehyde are potential inhibitors of QS.

Overall, the molecular dynamics analyses indicate that curcumin interacts more favorably with the quorum-sensing proteins than 5-bromoindole-3-carboxaldehyde. Curcumin forms a larger and more dynamic hydrogen-bond network, particularly with LasR and the PQS proteins, which contributes to stronger binding. In contrast, 5-bromoindole-3-carboxaldehyde forms fewer hydrogen bonds and exhibits a more rigid, protein-dependent binding mode, with reduced stability especially in the LasR complex.

These observations are supported by structural analysis based on RMSD and RMSF, which indicate that curcumin maintains stable binding throughout the simulations, reduces protein fluctuations and increases the dynamic stability of the complexes. Conversely, 5-bromoindole-3-carboxaldehyde induces higher flexibility in specific regions. Consistently, MM-PBSA calculations reveal that although all complexes are thermodynamically favorable, curcumin exhibits stronger van der Waals and electrostatic interactions, resulting in more favorable binding free energies across the three *P. aeruginosa* quorum-sensing proteins.

## Conclusion

This study provides strong evidence that both 5-bromoindole-3-carboxaldehyde and curcumin are effective in inhibiting quorum-sensing-dependent virulence factors in *Pseudomonas aeruginosa* in reference strains such as Pa01, Pa14 and ATCC 27853.

5-bromoindole-3-carboxaldehyde showed significant inhibition of pyocyanin, elastase, protease and alginate production, particularly in strain Pa01, which showed the greatest reductions in the levels of these virulence factors. The inhibition observed in strains Pa14 and ATCC 27853, although notable, was lower, suggesting further studies to elucidate the mechanism.

On the other hand, curcumin is a potent inhibitor of the same virulence factors in all strains studied, with a complete inhibition of pyocyanin production in strain ATCC 27853 at a concentration of 5 µg/ml. However, the efficacy of curcumin is variable between strains, with strain Pa14 showing the lowest overall sensitivity to this compound.

These findings support the hypothesis that inhibition of the quorum sensing system may be an effective strategy to reduce the virulence of *P. aeruginosa* without affecting its growth. The variability observed in the response of the different strains underlines the importance of considering bacterial heterogeneity when developing therapies based on quorum sensing inhibition.

## Acknowledgment

The researchers thank the academic direction of the degree in Biological Pharmaceutical Chemistry for support in carrying out this research work and Cristian Sadalis Santos López for their technical support.

## Code of ethics

In the present study, reference strains of *Pseudomonas aeruginosa* (Pa01, Pa14 and ATCC 27853) were used to investigate the quorum sensing inhibitory activity of curcumin and 5-bromoindole-3-carboxaldehyde. All experimental activities were carried out in compliance with BSL-1 level biosafety regulations, strictly following institutional protocols for the safe handling of microorganisms. The use of reference strains in this study does not involve humans or animals, and the samples used are of non-clinical origin and have been widely characterized and documented in the scientific literature. Therefore, ethics committee approval is not required for the performance of this work.

